# Modeling the photosynthetic system I as complex interacting network

**DOI:** 10.1101/2020.05.02.074377

**Authors:** D. Montepietra, M. Bellingeri, F. Scotognella, D. Cassi

## Abstract

In this paper we model the excitation energy transfer (EET) of the photosynthetic system I (PSI) of the common pea plant *Pisum Sativum* as complex interacting network. The magnitude of the link energy transfer between nodes-chromophores is computed by Forster Resonant Energy Transfer (FRET) using the pairwise physical distances between chromophores from the PDB (Protein Data Bank). We measure the global PSI network EET efficiency adopting well-known network theory indicators: the network efficiency (*Eff*) and the largest connected component (*LCC*). We find that when progressively removing the weak links of lower EET, the network efficiency (*Eff*) decreases while the EET paths integrity (*LCC*) is still preserved. This finding would show that the PSI is a resilient system owning a large window of functioning feasibility and it is completely impaired only when removing most of the network links. Furthermore, we perform nodes removal simulations to understand how the nodes-chromophores malfunctioning may affect the PSI functioning. We discover that the removal of the core chlorophylls triggers the fastest decrease in the network efficiency (*Eff*), unveiling them as the key component boosting the high EET efficiency. Our outcomes open new perspectives of research, such comparing the PSI energy transfer efficiency of different natural and agricultural plant species and investigating the light-harvesting mechanisms of artificial photosynthesis both in plant agriculture and in the field of solar energy applications.

## 1. INTRODUCTION

The ability of photosynthetic oxygenic organisms to convert light energy into chemical energy depends on a group of large membrane-bound complexes whose coordinated activity allows the capture of photons and their conversion into highly energetic molecules as NADPH and ATP through an electron transport chain [1]. The whole photosynthetic process is driven by the biological complexes possessing the light-capturing function: photosystem I (PSI) and photosystem II (PSII) [2,3]. The photosynthetic process starts with the catch of an incoming photon by PSII that utilizes it to oxidize water. The oxidation of water produces oxygen and reduces membrane-embedded quinones. The reduced quinones are then utilized by the cytochrome b6f complex to create a proton gradient across the membrane and to reduce the small copper protein plastocyanin (PC), that is the electron donor of PSI. After an additional photon is absorbed by PSI, an oxidation takes place and the removed electron migrates to finally reduce ferredoxin (Fd). Eventually, reduced Fd will reduce NADP to NADPH that will power the Calvin cycle to produce carbohydrates [4]. The photosynthetic process is schematically depicted in Figure 1.

**Figure 1:**
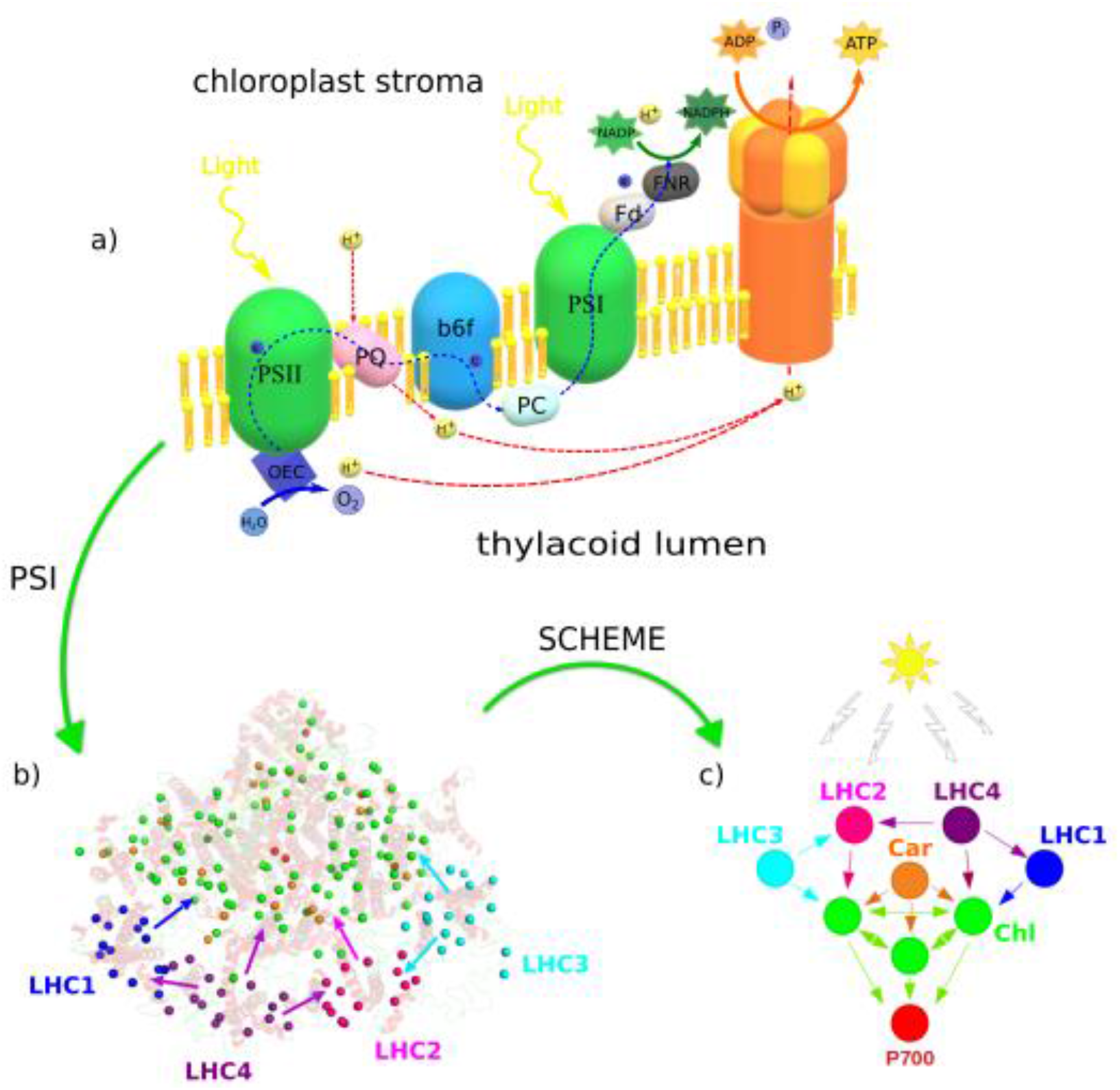
*a)* Schematic representation of the different protein complexes and the mechanisms involved in the photosynthetic process. In order, from left to right we find: OEC (Oxigen Evolving Complex), Photosystem II (PSII), Plastoquinone pool (PQ), b6f cytochrome (b6f), Plastocyanin (PC), Photosystem I (PSI), Ferrodoxin (Fd), Ferrodoxin NADP (+) Reductase (FNR) and ATP-synthase. *b)* Representation of the PSI network nodes embedded in the 3D protein structure. Nodes color indicates the different PSI network chromophores as in the right figure scheme (b). The arrows show the links between LHCs and other Chls with their directionality. *c)* Schematic representation of the different nodes of the PSI network and their connectivity. Car (orange) can only be starting nodes, thus they have only outgoing edges. We stress that every node can be the starting point of EET, but Car can only have outgoing links because of their higher site energy. Core Chls (green) can have both outgoing and ingoing edges. On the other hand, Chls belonging to different LHCs (LHC1 in blue, LHC2 in magenta, LHC3 in cyan and LHC4 in purple) can only trasfer energy within the same LHC or to Chls with lower energies. Finally, P700 (red) is the point of arrival of the FRET process and only possesses ingoing edges. The two P700 chlorophylls are not directly connected.

The light-harvesting biological complexes PSI and PSII are thought to be the fundamental (and probably the oldest) photosynthetic units [5]. Within the PSII and PSI complexes light is absorbed and transferred to a reaction center (RC) through a highly efficient Excitation Energy Transfer (EET) [6]. The two photosystems display a common structural organization and two main functional moieties: a core complex, containing the RC where the photochemical reactions occur, and a peripheral antenna system devoted to the increase of the light harvesting capability and to the regulation of the photosynthetic process [4,7,8]

The core complexes have been well conserved during the evolution, as most of the subunits are similar in prokaryotic and eukaryotic photosystems and only a few are specific to each group. On the contrary, the peripheral antenna system displays great variability, being composed of peripheral associated membrane proteins in cyanobacteria, called phycobilisomes, and integral Light Harvesting Complex (LHC) membrane proteins in eukaryotic cells [9].

The light-harvesting process and the EET from the antenna complexes to the RC are facilitated by pigment-protein complexes (PPCs). In green plants (Viridiplantae), most of the light is absorbed by the photosynthetic pigments chlorophyll *a* and *b*, allowing efficient light harvesting and ultrafast EET amongst antenna chlorophylls, leading to the quantum and thermodynamic efficiencies which are the highest known [9,10]

When a photon is absorbed, a chlorophyll (Chl) is excited to the singlet excited state (1Chl*). In PSI this excitation energy is transferred from the antenna pigments to the primary electron donor P700, a special Chls pair located at the RC. The excitation energy is used to build up the singlet excited state P700*, leading to charge separation, where P700^+^ remains and the electron is transferred across the thylakoid membrane by a chain of electron carriers.

Other cofactors present in PSI are also involved in the processes of light capturing, charge separation and prevention of photodamage, like carotenoids (Cars). This network of pigments, whose absorption spectra spans a broad spectral range, is embedded in a protein scaffold which enforces chromophores spatial organization by holding them in a relatively rigid position and orientation [3].

While huge amount of research has been focused at understanding the PSI core electron transfer chemistry which is induced by the arrival of excitation energy at the RC [11–17], few attention has been directed to unveil the underneath structural and topological patterns shaping the high EET efficiency of the PSI system [2,3].

The purpose of this research is the analysis of the structural organization of a plant PSI as a complex networked system for EET by means of network science viewpoint. Network science has proven to be a powerful tool for the study of complex systems from many different realms [18–26]. Basically, a network is a model composed by nodes (or vertices) and links (edges), where nodes indicate the components of the system and the links define the interactions among them [27,28]. Here we model the PSI light-harvesting system of the plant *Pisum sativum* as a complex network of nodes-chromophores describing the energy transfer links among them. First, we find that the PSI is a highly connected network with very short EET pathways from the nodes harvesting the light to the RC. Second, we discover that the capacity to perform EET in the whole network is severely impaired only by removing the major amount of EET links. This unveils the high resilience of the PSI system, which holds the capacity to perform the energy transfer in the whole network even reaching severe conditions for its feasibility. Last, we simulate different scenarios of chromophores malfunctioning by removing nodes from the network [18,29]. We find that the removal towards the core Chls is clearly more harmful in decreasing the PSI functioning, suggesting the core as the key structural component of the high efficiency EET in the PSI of the *P. Sativum*.

## 2. METHODS

### 2.1 Network theory

A generic network (or graph) G(*N,L*) is composed by a set of *N* nodes and *L* links connecting them. The network G can be abstracted in *N*x*N* square adjacency matrix A, where the element a_*i.j*_ =1 indicates the presence of a link between node *i* and node *j* and 0 otherwise [27]. In the case G is undirected, e.g. the link have no directionality, the adjacency matrix is symmetrical with a_*i.j*_ In a directed network the adjacency matrix is not symmetrical, the links own proper direction and a_*i.j*_ =1 stands for a link from *i* to *j*, and a_*j.i*_ = 1 indicates that the link goes from *j* to *i*. In the case the links among nodes are only present-absent, as in G above described, the network is called binary or topological or unweighted. Nevertheless, many real networks are naturally weighted displaying a large heterogeneity in the capacity of the links-connections. In fact, in weighted network links are associated with weight values that differentiate them in terms of their strength, intensity, or capacity [19,22,30–33]. Thus, a weighted network can be abstracted by a *N*x*N* weighted adjacency matrix W, where the element *w_i,j_* > 0 in the case there is link of weight *w_i.j_* between the node *i* and the node *j* and *w_i.j_* = 0 otherwise. Many empirical researches demonstrate that purely binary-topological models are inadequate to explain the complex structure of the real systems, and that there is a need for weighted network models to overcome the pure topological description [27,28,34]

### 2.2 Building the Photosystem I Network (PSI)

We build the PSI network of the common pea plant (*P. sativum*) starting from the PDB (Protein Data Bank) entry 4RKU. In the PDB, we find all available crystals of PSI belong to this species. We model the PSI as weighted directed network of *N* nodes-chromophores and *L* links-EET. The nodes represent chromophores with different functions, i.e. β-carotenes (Cars) and chlorophylls (Chls a or b). To model the PSI network we assumed that Cars and Chls can be summarized by one of their atoms. In fact, Cars are long conjugated molecules with electrons delocalized in the entirety of the molecule, making the internal processes much faster than the external ones (i.e. energy transfer) [35]; so each Car can be considered almost like a point-node object delivering energy in the system. For this reason, we choose the C15 carbon, located in the middle of the molecule, to model the carotenoids. We found 21 Cars molecules in the structure. For Chls we represented the molecule using its Mg atom, that is a common practice in the literature [6,36]. We found 158 Chl molecules in the PSI complex (including P700). In this way every node of the PSI network is a C or a Mg atom, representing the corresponding molecule. Modeling the PSI we obtain a network with a total number of nodes N = 179.

Then, we assume that the underlying mechanism of energy transfer (EET) between Chls, and between Cars and Chls is represented by Forster Resonant Energy Transfer (FRET) [37]. FRET contributes to the most of the EET acting as a good approximation to investigate energetic coupling of the underlying spatial organization of the chromophores [38,39]. Future investigations may expand the model presented here by including the quantum mechanical dynamics to model the EET of the PSI.

The FRET efficiency (*E*):

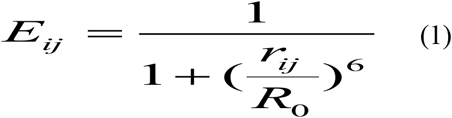

where E_*ij*_ is the FRET efficiency the link connecting nodes *i* and *j*, *R*_0_ is the Forster distance between *i* and *j*, and *r_i,j_* is the physical distance between the nodes. All pairwise physical distances between nodes can be calculated directly from their PDB spatial coordinates. Therefore, *E_ij_* represents the weight *w_ij_* of the link joining node *i* and node *j*. The value of *R_0_* of Chls-Chls FRET is 90 Å [40,41], and we assume the same value for Cars-Chls FRET. Furthermore, in the photosynthetic process, Cars absorb and transfer lower wavelength photons to Chls, which in turn cannot transfer the excitation back to Cars but only send it to other Chls, until they reach the P700 [42]. For this reason, carotenoids will be represented by nodes with outgoing links only. On the contrary, the two Chls forming P700 will have only incoming links as they can only receive excitation; furthermore, the Chls of the P700 do not perform EET and thus no direct link connection is present between them. Last, with the sake of simplicity, our model does not consider the competition between parallel energy transfers that may induce radiative (light emission) and non-radiative (heat production) de-excitation, with a global EET efficiency decreasing.

### 2.3 The PSI network with the energy funnel

Light-harvesting complexes (LHCs) chlorophylls are the light entry gate of photosynthesis [43]. The different LHCs Chls are used to absorb light in spectral regions where more radiant energy is available, increasing the amount of sunlight that can be collected [44]. In the PSI, four types of Chls LHCs are found: LHC1, LHC2, LHC3, LHC4 [4]. Their schematic representation can be seen in Fig. 1. Different LHCs have different light absorption peaks, corresponding to different site energies of the Chls. A photon with a certain energy absorbed in one of these LHC will therefore be directed only to Chls with lower or equal site energies. This generates an energetic funnel, that it is used by the photosynthetic organisms to speed up and increase the EET efficiency [45]. The first step in modelling this funnel is thus the assignment of the different site energies to the LHCs, that will define the connectivity among them. We assign the site energies of the LHCs as in [38] (Table 2). Further, [38] showed that inter-LHCs EET is much less efficient than LHC-core EET. Following this finding, we assume that only adjacent LHCs can transfer energy between them. We depict the possible links among LHCs complexes in Fig. 2. In Table 1 we summarize the different nodes-chromophores and their properties.

**Figure 2:**
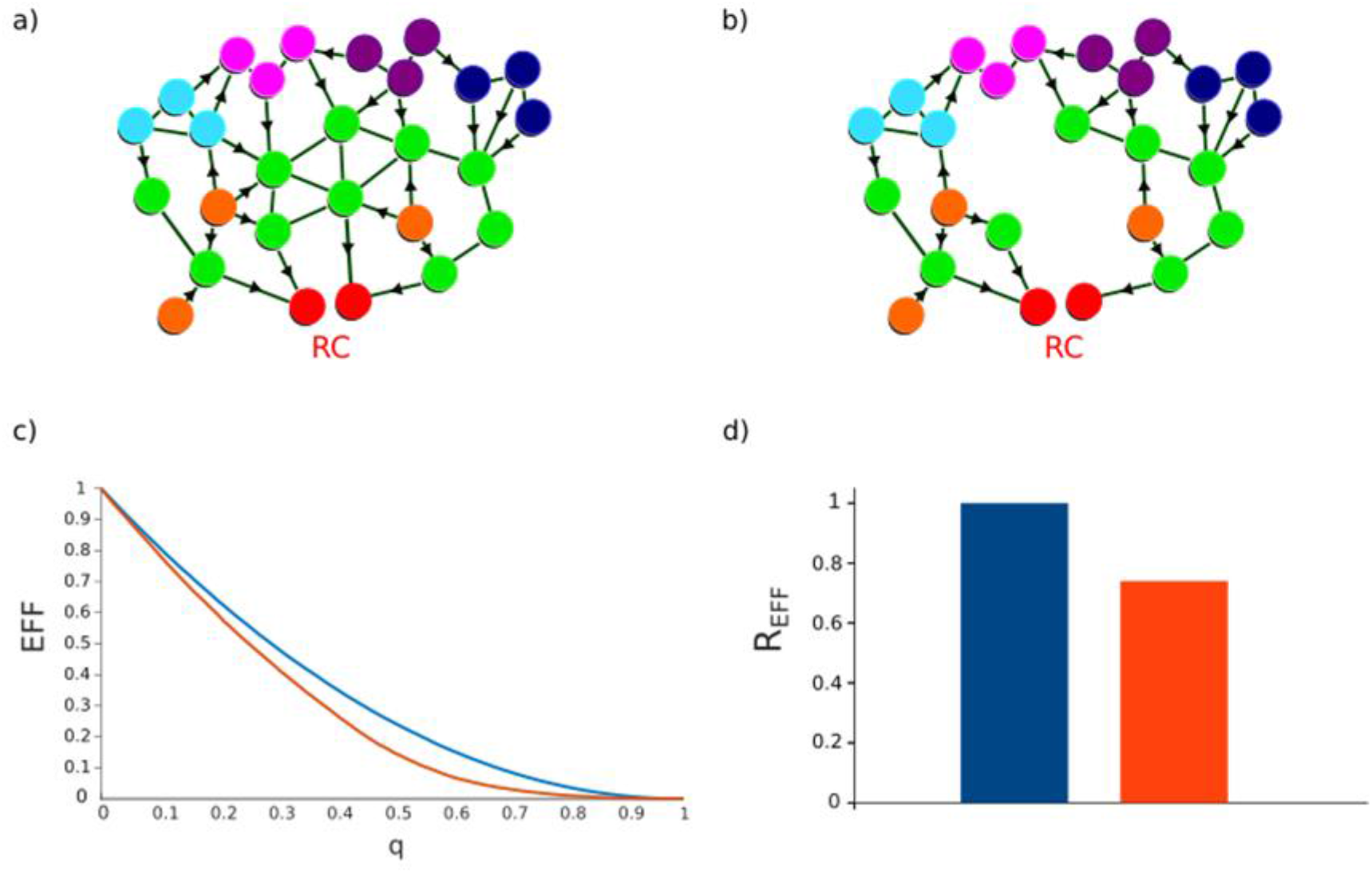
example network where nodes-chromophores perform EET toward the RC (a); the network in (a) after the removal two green nodes, where we can see the elongation of the EET paths and the consequent network functioning decrease; functioning efficiency measure (*Eff*) as a function of the fraction of nodes removed (*q*) for two hypothetical different attack strategies (c); the red strategy triggers a faster efficiency (*Eff*) decrease than the blue strategy and the network robustness area below the red curve is lower than the one below the blue curve. The robustness (*R_Eff_*) of the efficiency computed for the two strategies (d). The bar robustness *R_Eff_* corresponds to the area below the curve in the left chart (c); *R_Eff_* value is normalized by the max value, e.g. by *R_Eff_* of the blue strategy.

**Table 1:** Summary of the different nodes-chromophores of *P. Sativum* PSI network.

| Node Type | Acronyms | Number | Biological function | Link direction |
| --- | --- | --- | --- | --- |
| Carotenoids | Cars | 21 | Photodamage protection;<br>Lower wavelength absorption;<br>Energy transfer | Outgoing |
| Core Chlorophylls | Core Chls | 97 | Energy transfer | Outgoing + ingoing |
| Ligh Harvesting<br>Complexes<br>Chlorophylls | LHCs | 59 | Light harvesting;<br>Wider harvested light spectrum;<br>Energy transfer | Outgoing + ingoing |
| P700 | P700 | 2 | Reaction center of PSI, special<br>pair of chlorophylls | Ingoing |

**Table 2:**
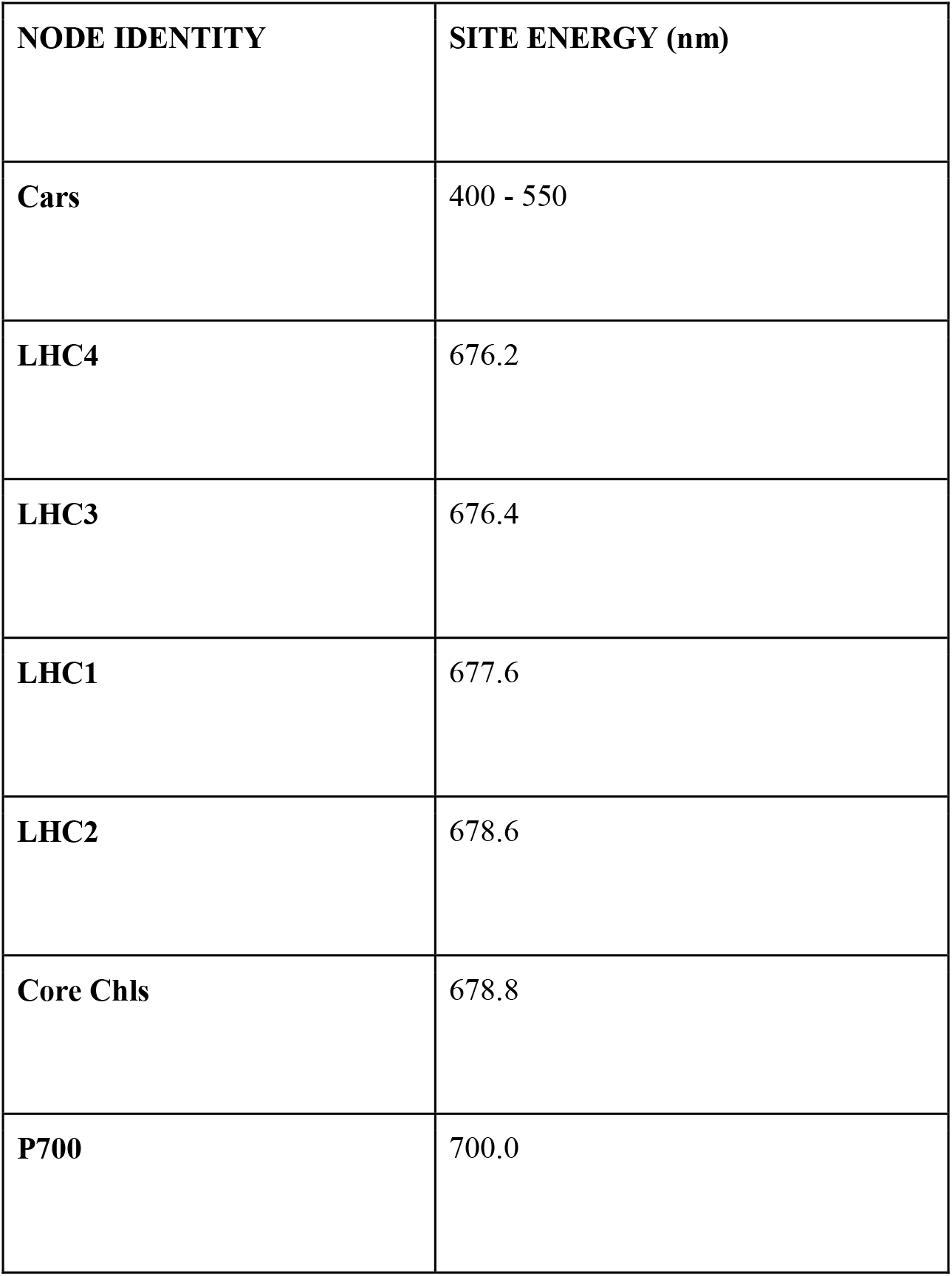
Site energies related to different types of node-chromophores.

### 2.4 The cut-off distance (CD)

The links of the PSI network own different FRET efficiencies as a function of the nodes pairwise distances of the PDB structure. The closer the nodes, the higher the link weight (FRET efficiency) joining them. For this reason, we can build the PSI network with different number of links as a function of their FRET magnitude. To do this, we define the cut-off distance (CD) as the distance between two nodes over which we assume the link between them absent because the associated FRET efficiency computed by Eq. 1 is too low. Thus, by lowering the CD parameter, we require shorter pairwise distance to form links among nodes. In other terms, starting from the network where all links are drawn, when lowering the CD, we operate a removal of the links of lower weight (weak links) maintaining the links of higher weight (strong links). We created several networks using the cut-off distance values: NO cut-off, 90, 80, 70, 60, 50, 40, 30, 20 and 10 Å.

### 2.4 Measures of network functioning

#### The largest connected component (*LCC*)

The *LCC* is a widely used measure of the network functioning [18,46,47]. The *LCC* is also known as giant component and it accounts the highest number of connected nodes in the network. We compute two types of *LCC*, by considering the weakly largest connected component (*LCC_weak_*) and strongly largest connected component (*LCC_strong_*). The *LCC_weak_* represents the largest number of nodes in which every node can reach any other by an undirected path. Differently, the *LCC_strong_* represents the largest number of nodes in which every node can reach any other node by a directed path [27].

#### The network efficiency (*Eff*)

*Eff* is a widely used measure of network functioning evaluating how efficiently the network exchanges information [20,48,49]. *Eff* is based on the shortest paths (SP) notion. A path is a sequence of links connecting two nodes in the network; a binary shortest path between a pair of nodes is the minimum number of links that we have to pass travelling between the nodes [50]. Differently, the weighted shortest paths (wSP) consider both the number of links and the weight attached to them. To compute wSP, we first compute the inverse of the link weights; then we compute the weighted shortest paths as the minimum sum of the inverse link weights necessary to travel among nodes [27,34].

The average efficiency of the network is then defined:

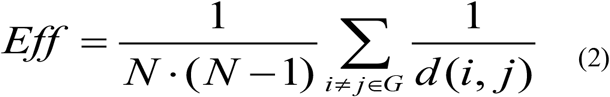

where *d_i,j_* is the shortest path between node *i* and node *j*, and *N* is the total number of network nodes. Eq. 1 indicates that the longer the shortest paths among nodes, the lower the efficiency of the network. The efficiency can be computed for undirected and directed networks when considering undirected and directed shortest paths respectively. Furthermore, the network efficiency can properly evaluate binary and weighted networks; in the case the network is weighted, *d_i,j_* indicates the weighted shortest path between nodes *i* and *j*. Since the PSI is a weighted and directed network, we use the weighted and directed network efficiency (*Eff*).

#### The number of connected nodes to P700 (*CN*)

We account the number of connected nodes to P700 through a directed path. This measure indicates how many nodes in the network can actually transfer energy (directly or indirectly) to the P700 reaction center. In other words, *CN* accounts the number of nodes-chromophores that can properly contribute to the PSI functioning.

### 2.5 The PSI network properties

We analysed the PSI system using well known notions from networks science. **Nodes degree (*k*)**: is the total number of links to the nodes [27]. We also computed the number of ingoing links, i.e. the indegree (*k_in_*) and the number of outgoing links, the out-degree (*k_out_*). **Nodes Strength (*S*)**: is the sum of the link weights to the nodes [49]. The strength of a node is the weighted counterpart of the node degree (*k*). Analogously we can compute the ingoing strength (*S_in_*) and the outgoing strength (*S_out_*). **Nodes betwenness centrality (*BC*)**: is a widely used measure of nodes centrality based on shortest paths. The *BC* of a node is the number of the shortest paths between all nodes pairs that pass through that node [27,51]. Here we adopt both the binary betwenness centrality (*BC*) and the weighted version of the betwenness centrality of the nodes (*BCw*). The *BCw* is computed using the weighted shortest paths. The weighted betweenness centrality of a node accounts the number of weighted shortest paths from any couple of nodes passing along that node. The higher is the *BCw* of a node, the higher is the number of weighted shortest paths passing along it (and the more central is the node). **δ centrality**: the node δ centrality accounts the decrease of the network efficiency (*Eff*) triggered by the removal of that node from the network [48] The higher the δ, the higher the contribution of the node on the network efficiency (*Eff*).

### 2.6 The nodes removal simulations

We simulated different nodes removal (attack) processes over the PSI network [18,52]. The PSI network nodes removal may describe functionality loss of the chromophores, that can be due by aging, disease, pollution or others [3].

The nodes removal strategies are:

- **Rand:** network nodes are randomly removed. It represents the case of failures in the network and it is used as benchmark comparison of the harmful of the selective attack strategies [18,29].
- **Rand LHC:** in this strategy we randomly remove nodes in the LHC ensemble that is composed by 59 nodes.
- **Rand Core:** in this strategy we randomly removed nodes in the Core Chls ensemble that is composed by 97 nodes.
- **Rand Car:** in this strategy we randomly removed nodes in the Cars ensemble that is composed by 21 nodes.

We measured the functioning of the network during the nodes removal process using the network directed weighted efficiency (*Eff*) and the weakly largest connected component (*LCC_weak_*). Since the random removals are stochastic processes, we average the outcomes from 10^3^ simulations. The damage triggered in the system by the specific PSI nodes removal (Rand Car, Core and LHC) is then compared with the damage induced by the random removal (Rand) of the same number of nodes. In this way we can discriminate the importance of the different types of nodes-chromophores.

### 2.7 The robustness of the network

When nodes are removed from the PSI network, we assist to the elongation of the EET paths and consequent system functioning decrease (Fig. 2 a, b). We can assess the system damage subjected to nodes removal according to the functioning measures. The more important the nodes removed from the network, the steeper is the decrease in the network functioning measure. For example, in the Fig. 2c the red removal strategy identifies more important nodes causing a steeper decrease in the network functioning efficiency (*Eff*) with respect to the blue strategy. To compare the response among nodes removal strategies, we resume the outcome in a single value defined network robustness (*R*), depicted by the bar plots of the Fig. 3d. The value of *R* corresponds to the area below the curve of the system functioning measure subjected to nodes removal. We referred to *R_EFF_* and *R_LCC_* when the network functioning is evaluated by the *Eff* and *LCC* respectively.

**Figure 3:**
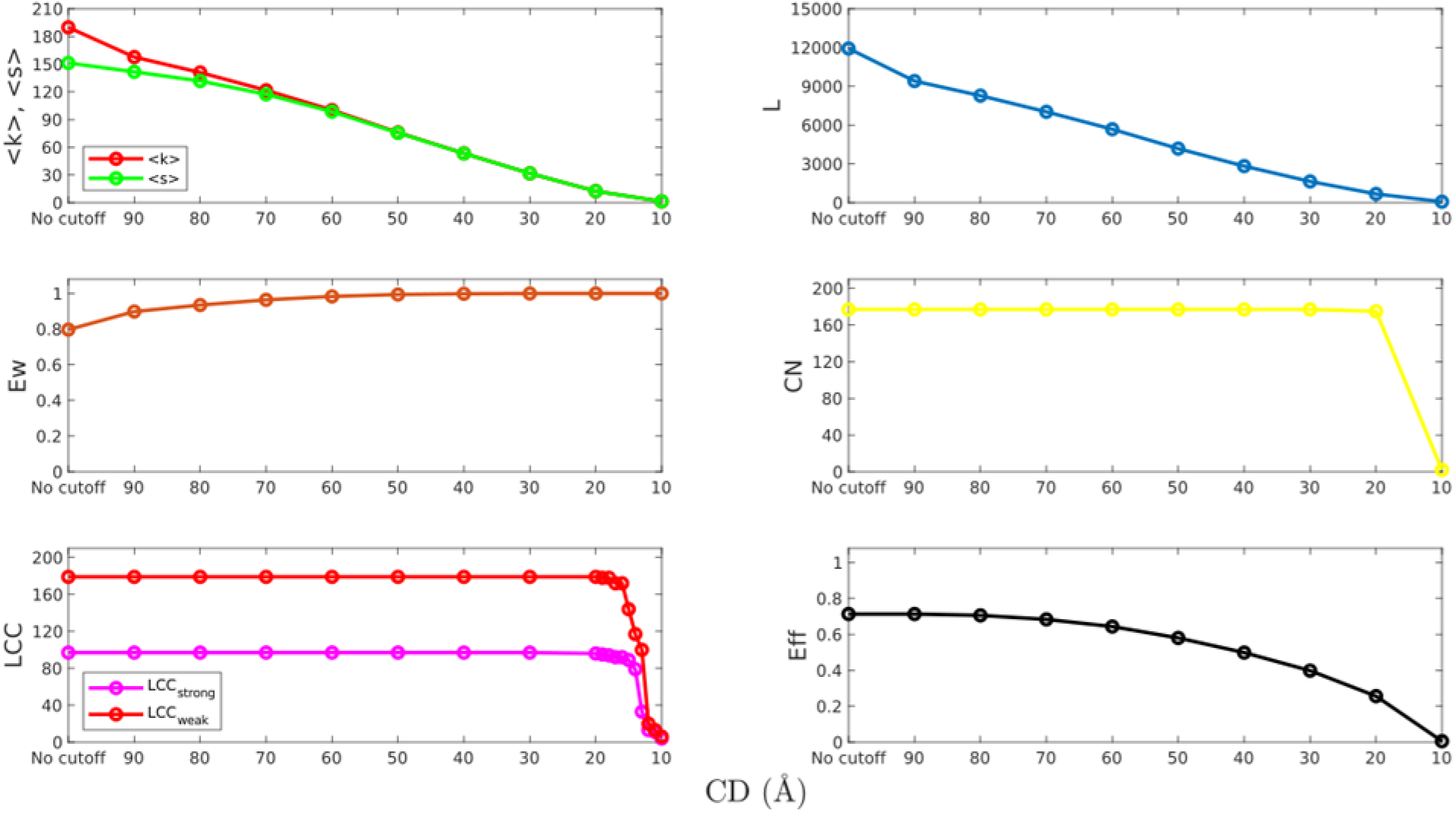
Network properties as a function of CD. average nodes degree <*k*>, average nodes strength <*s*>, largest connected component *LCC*, number of links *L*, network efficiency *Eff*, average link weight *Ew, CN* number of connected nodes to the P700.

## 3. RESULTS

### 3.1 The network properties

In Table 3 we resume the PSI network properties for each CD value. The number of links *L*, average nodes degree <*k*> and average nodes strength <*S*> linearly decrease lowering CD (Fig. 3). The nodes degree *k* ranges in the interval 0<*k*<270. The nodes degree distribution is: a tri-modal with three bell-shaped distributions from NO CD to CD=50 Å, one bell-shaped distribution with most of the values approaching the mean from CD=40 Å to CD=20 Å, and it becomes a Dirac distribution where all nodes present degree 1 for CD=10 Å (Appendix, Fig. A1). The node strength distribution follows the node degree distribution patterns indicating a nodes degree-strength coupling (Fig. A2). The average link weight <*Ew*> shows a very slow increase lowering CD (Fig. 3); the link weight distributions are highly skewed to the left with many links presenting weight 1 and few links of lower weight (Fig. A3). The network efficiency (*Eff*) remains roughly constant by decreasing CD from NO CD to CD=70 Å; after this value, we find a more pronounced *Eff* decrease reaching the minimum for CD=10 Å (Fig. 3). *LCC_weak_* spans the whole network with *N*=179 nodes up to reach CD>10 Å (Fig. 4). *LCC_strong_* holds the values *N*=97 up to reach CD>10 Å (Fig. 3). We find 177 nodes connected to the P700 (*CN*) and this number holds constant up to reach CD=20 Å (Fig. 3). Only for CD=10 Å *CN* drop to *quasi* zero. It is worth noting that for *LCC_weak_, LCC_strong_* and *CN* measures the critical CD value is 10 Å, below which the measures quickly collapses.

**Figure 4:**
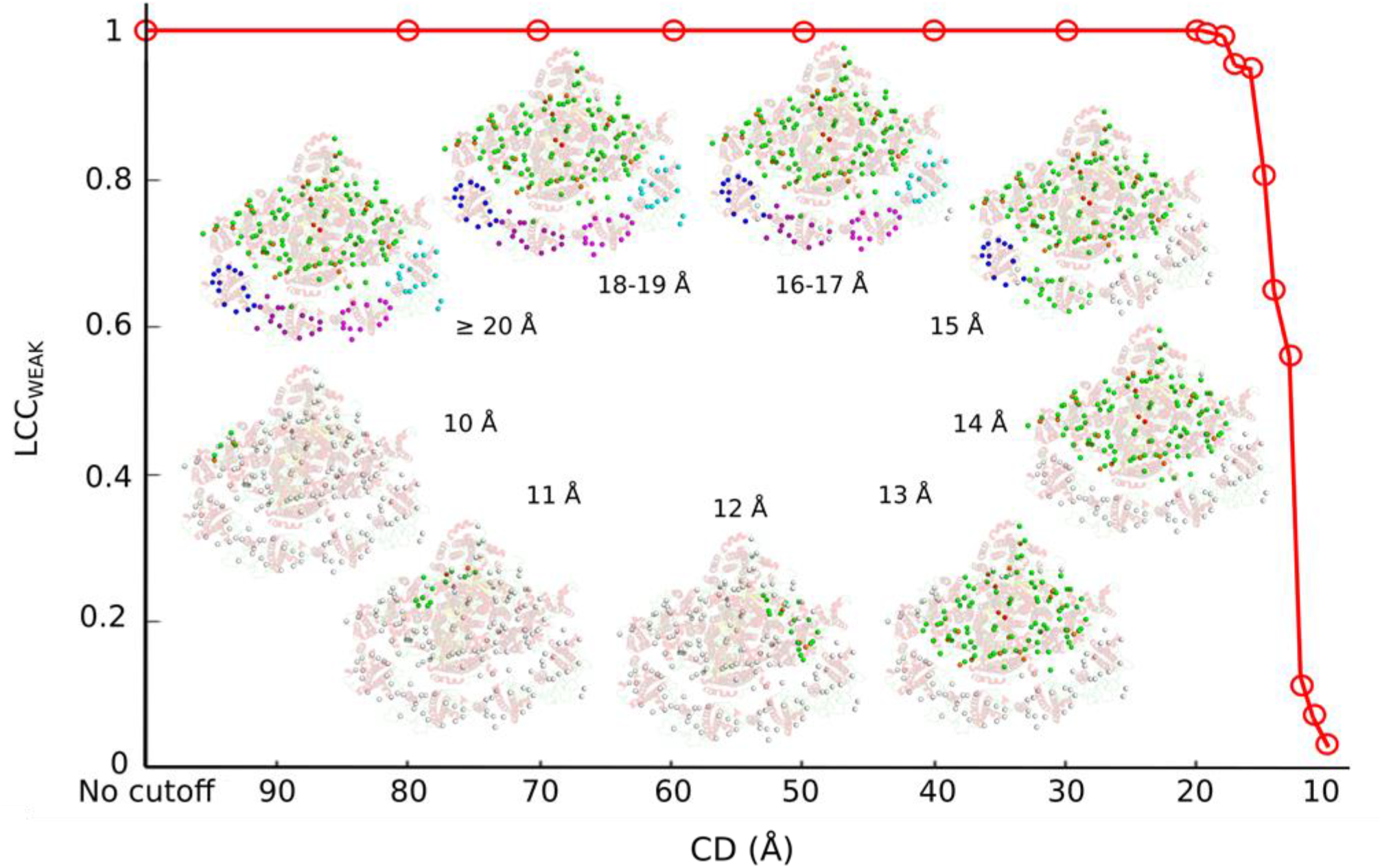
The normalized *LCC_weak_* size and the representation of the various *LCC_weak_* of the PSI network found at different cut-off distances (CD) as seen in the underlying protein structure. The nodes are colored while they appear in the *LCC* according to the scheme colors used in Fig.1.

**Table 3:** PSI network features for each CD value. <*k*> average nodes degree, <*k_out_*> average nodes out-degree and <*k_in_*> average nodes in-degree; *L* total number of links; <*S*> average node strength, <*S_in_*> average in-strength, <*S_out_*> average out-strength; *Ew* average link weights; *Eff* global network efficiency; *LCC_weak_* weakly largest connected component; *LCC_strong_* strongly largest connected component; *CN* number of connected nodes to the P700.

| CD | $\langle k \rangle$ | $\langle k_{out} \rangle$ | $\langle k_{in} \rangle$ | $L$ | $\langle S \rangle$ | $\langle S_{out} \rangle$ | $\langle S_{in} \rangle$ | $Ew$ | $Eff$ | $LCC_{weak}$ | $LCC_{strong}$ | $CN$ |
| --- | --- | --- | --- | --- | --- | --- | --- | --- | --- | --- | --- | --- |
| NO | 190 | 95 | 95 | 11924 | 151.3 | 75.65 | 75.65 | 0.7972 | 0.71 | 179 | 97 | 177 |
| 90 Å | 158 | 79 | 79 | 9409 | 141.68 | 70.84 | 70.84 | 0.8984 | 0.71 | 179 | 97 | 177 |
| 80 Å | 141 | 70 | 70 | 8279 | 131.79 | 65.9 | 65.9 | 0.9348 | 0.71 | 179 | 97 | 177 |
| 70 Å | 122 | 61 | 61 | 7028 | 117.32 | 58.66 | 58.66 | 0.9640 | 0.68 | 179 | 97 | 177 |
| 60 Å | 100 | 50 | 50 | 5681 | 98.55 | 49.28 | 49.28 | 0.9832 | 0.64 | 179 | 97 | 177 |
| 50 Å | 76 | 38 | 38 | 4181 | 75.54 | 37.77 | 37.77 | 0.9939 | 0.58 | 179 | 97 | 177 |
| 40 Å | 53 | 27 | 27 | 2828 | 53.3 | 26.65 | 26.65 | 0.9982 | 0.50 | 179 | 97 | 177 |
| 30 Å | 32 | 16 | 16 | 1642 | 31.74 | 15.87 | 15.87 | 0.9996 | 0.40 | 179 | 97 | 177 |
| 20 Å | 12 | 6 | 6 | 676 | 12.48 | 6.24 | 6.24 | 1.0000 | 0.26 | 179 | 97 | 175 |
| 10 Å | 1 | 1 | 1 | 74 | 1.39 | 0.69 | 0.69 | 1.0000 | 0.01 | 6 | 4 | 2 |

### 3.2 The properties and connectivity of the different nodes-chromophores

The Core Chls show the highest degree and in-degree (*k* and *k_in_*), betwenness centrality (*BC* and *BCw*), strength (*S* and *S_in_*) and δ centrality than all the others types of nodes-chromophores (Figs 5–6); the LHCs show the highest outgoing degree (*k_out_*) and strength (*S_out_*) (Figs 5–6); the P700 has the highest in-degree (*k_in_*, comparable to the Core) and in-strength (*S_in_*) (Figs 5–6). We find a higher number of LHCs-Core links than the number of intra-LHCs links, e.g. each LHC shares more than 1200 links with the Core Chls and *circa* 100 intra-LHCs links for CD=NO (Fig. 7). Nonetheless, when lowering CD, we find that the number LHCs-Core links tends to decrease, while the number of intra-LHCs links holds roughly constant. For all LHCs we find that the number of LHC-Core links become similar to the number of intra-LHCs links for CD=50 Å (Fig. 7).

**Figure 5:**
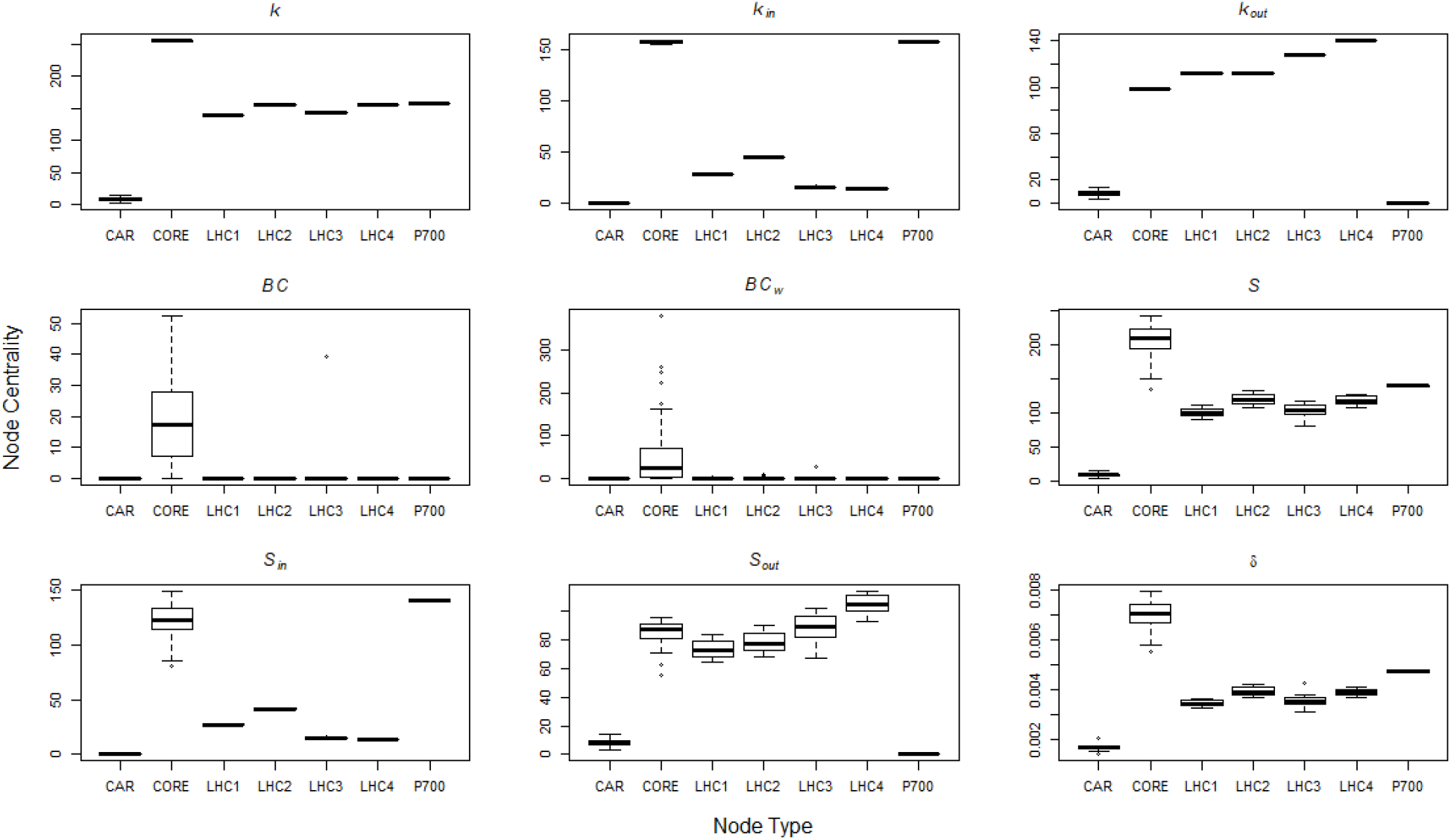
Node types *vs* nodes centrality feature for the PSI network features for no cut-off (CD=NO). Keys are: *k* degree of the nodes, *k_out_* out-degree, *k_in_* in-degree; *BC* binary directed betwennness centrality; *BCw* binary directed betwennness centrality; *S* node strength, *S_in_* ingoing node strength, *S_out_* outgoing node strength; δ decrease in the directed network efficiency after single node removal.

**Figure 6:**
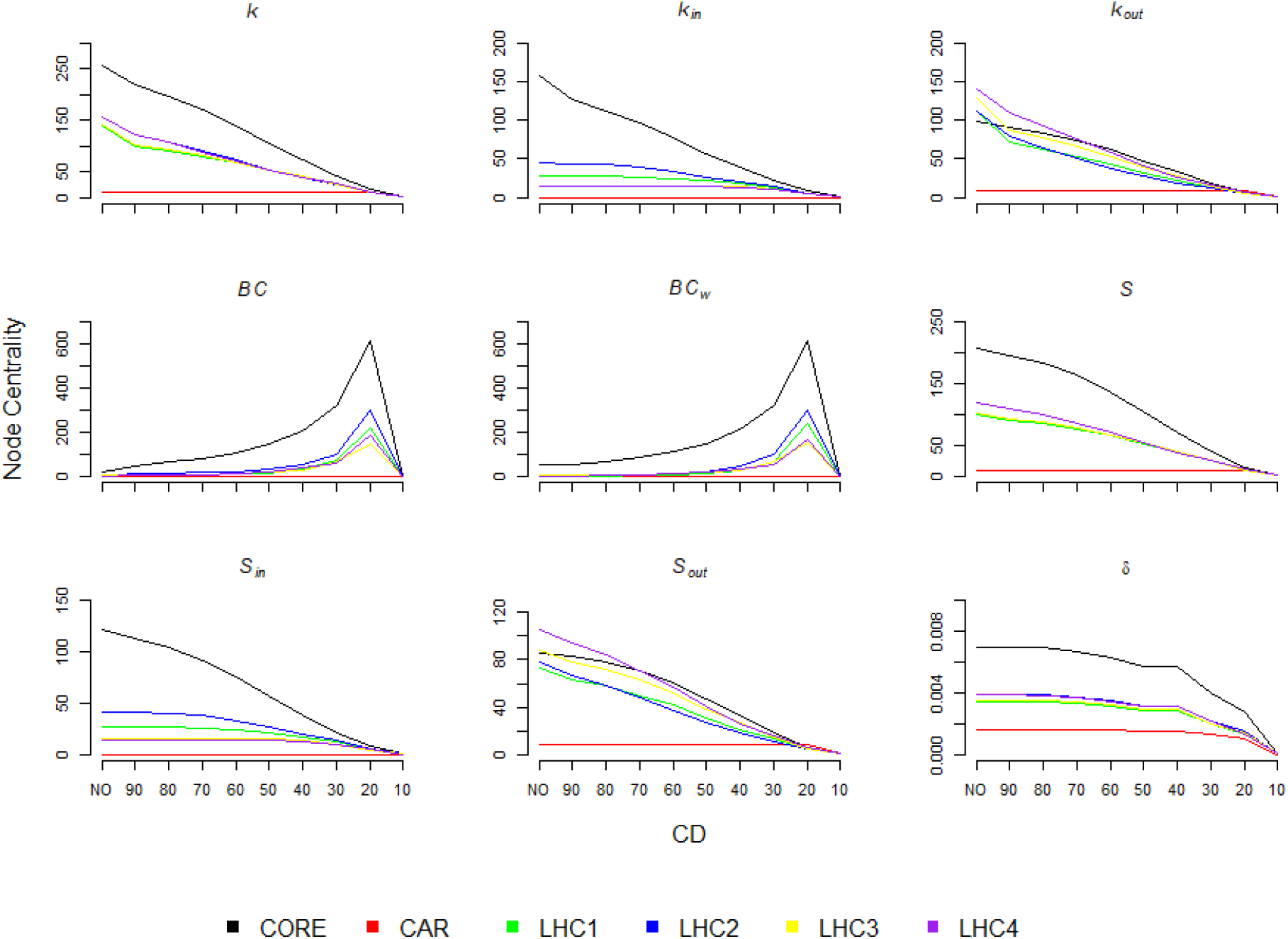
Node centrality features for the different types of PSI network chromophores as a function of the CD. Keys are: *k* degree of the nodes, *k_out_* out-degree, *k_in_* in-degree; *BC* binary directed betwennness centrality; *BCw* binary directed betwennness centrality; *S* node strength, *S_in_* ingoing node strength, *S_out_* outgoing node strength; δ decrease in the directed network efficiency after single node removal.

**Figure 7:**
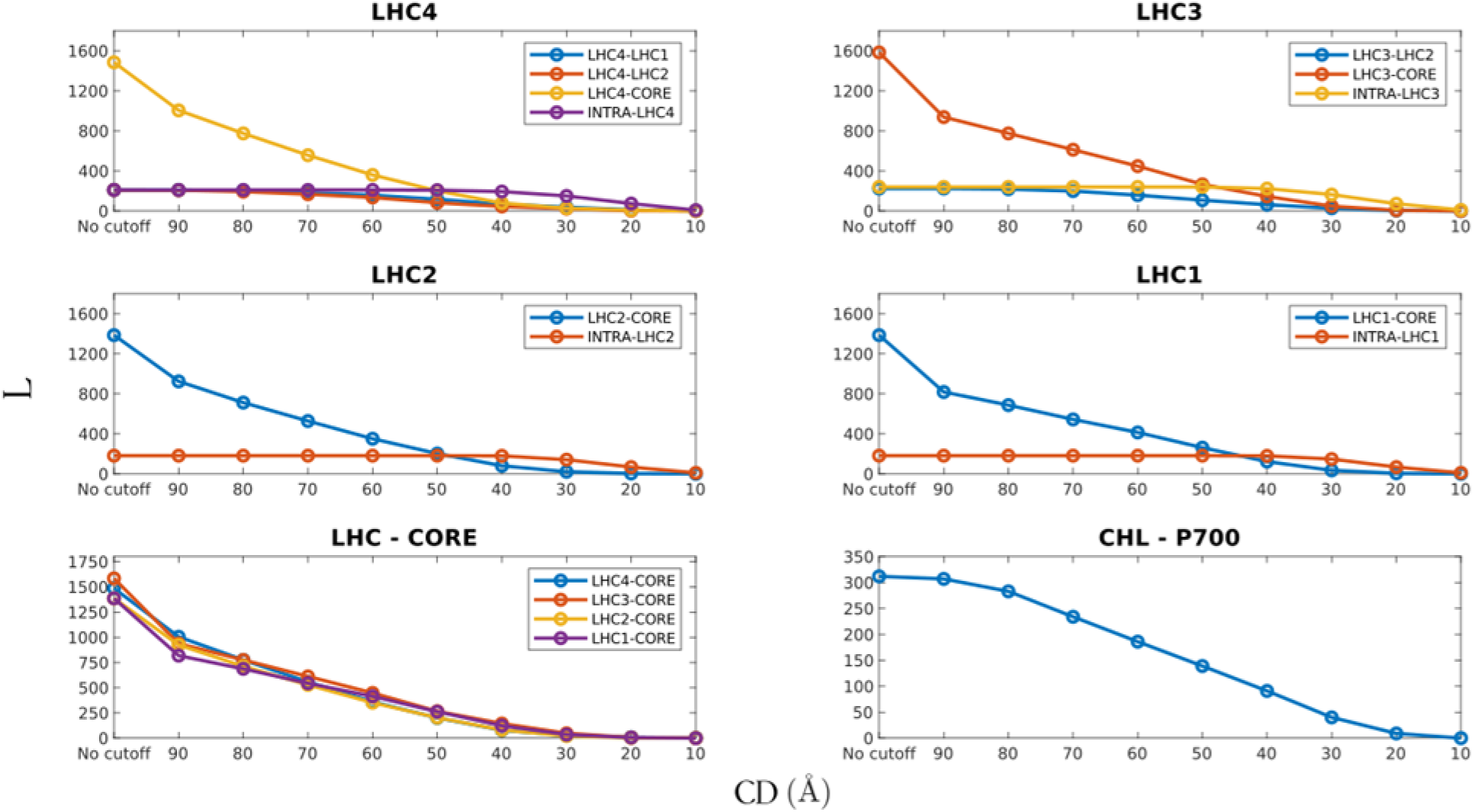
Number of links *L* among different types of nodes-chromophores as function of CD.

### 3.3 Nodes removal

#### Carotenoids

Rand Car induces less average damage to the *Eff* (higher *R_Eff_*) of the PSI network than the random removal control (Rand) (Fig. 8). The Rand Car shows roughly half of the *R_EFF_* (50% fewer) of Rand. This difference between the *R_Eff_* values decreases by lowering CD. Also with the *R_LCC_*, Rand Car induces less average damage to the *LCC_weak_* than Rand (Fig. A13). The difference between the *R_LCC_* values of the Rand Car and Rand is less pronounced for CD=10 Å (Fig. A13).

**Figure 8:**
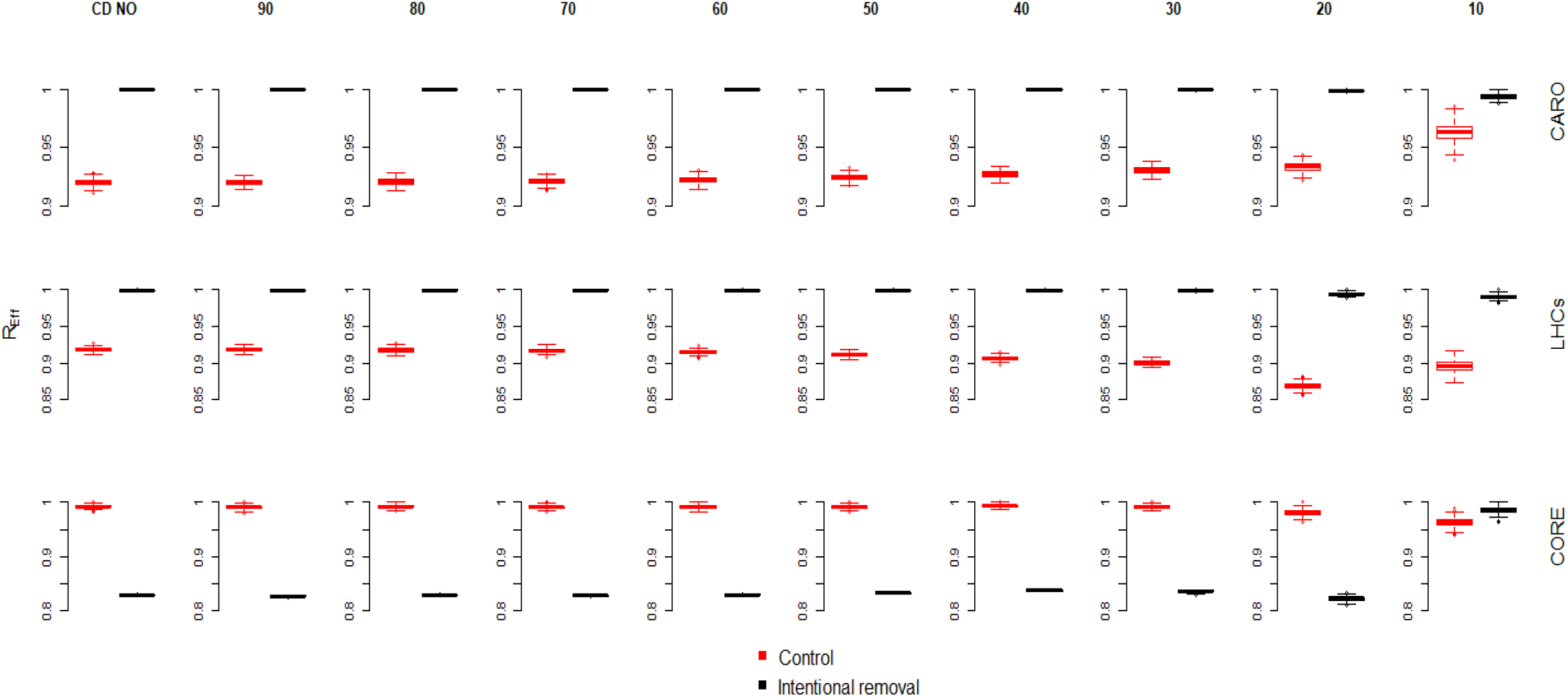
PSI network robustness (*R_Eff_*) under the random removal (Rand) and the intentional random nodes removal strategies for the different CD. The intentional random removals: top row Cars, half row LHCs and bottom row Core Chls.

#### LHCs

Random removal of LHCs Chls (Rand LHCs) causes less damage to *Eff* (higher *R_Eff_*) than the random removal control (Rand) for all CDs (Fig. 8). The Rand LHCs induces less average damage to the *LCC_weak_* (higher *R_LCC_*) of than Rand (Fig. A13). This difference is less pronounced for CD=10 Å (Fig. A13).

#### Core

Random removal of Core Chls (Rand Core) causes more damage to *Eff* (lower *R_EFF_*) than the random removal control (Rand) (Fig. 8). The *R_EFF_* obtained by Rand Core is *circa* 25% lower than the one induced by the Rand. Only CD = 10 Å, Rand Core triggers greater damage than Rand. Rand Core causes more damage to *LCC_weak_* than Rand (Fig. A13). This difference is less pronounced for CD=10 Å.

## 4. Discussion

### 4.1 Network properties

The Core Chls nodes show the highest degree (*k*) among all other types of chromophores, i.e. they share the higher number of links in the network (Figs 5–6) indicating the Core as the PSI network component playing a major role for the global energy transfer. This is clearly confirmed by their highest betwenness centrality for all CDs (Figs 5–6). The nodes with highest betwenness are the articulation points routing the shortest and the most efficient energy transfer paths in the network. As a consequence, these findings demonstrate that the Core is the ‘highly connected’ and ‘most central’ component in the PSI network of the *P. Sativum*, corroborating previous outcomes from literature showing that the primary function of the Core Chls is to enhance the EET efficiency in the system [6,53].

Differently, the discovery that the LHCs nodes own the highest number of outgoing links (*k_out_*) than all other types of nodes chromophores and the Cars nodes have only outgoing links (Figs 5–6) would indicate these types of chromophores as important light entry gate. This would confirm empirical evidences from literature, showing that Cars and LHCs have the primary function of light harvesting, and then perform energy transfer to other types of nodes throughout the outgoing links [6,35,53].

### 4.2 Feasibility of the PSI system

When lowering the cut-off distance (CD), shorter pairwise distance is required between nodes to perform FRET energy transfer links. In other terms, lowering the CD we progressively remove from the network the weak links of lower FRET efficiency and we maintain only the most efficient and most probable FRET links (strong links). The strong links connecting closest chromophores can be viewed as the ‘feasible FRET’ even when the system works at very low regime of functioning, such as scarce light radiation [54]. The low level of functioning may also occur after the vanishing of FRET links, such as for Chls ageing following the life of the leaf, with older PSI still possessing the higher FRET links only [55]. Therefore, CD tuning gives us information on the PSI system feasibility at different level of functioning efficiency and its resilience with regards of leaf ageing.

We discover that lowering the CD, the number of feasible energy transfer links (*L*) linearly decrease (Fig. 3). This decrease triggers the elongation of the EET paths in the PSI network and the consequent lowering of its efficiency *Eff* (Fig. 3). Hence the PSI system experiences a reduction in the network efficiency, the *LCC* representing the maximum number of connected nodes-chromophores holds constant up to reach CD=20 Å. Further, the number of nodes connected by directed paths to the P700 reaction centre (*CN*) hold constant up to reach CD=20 Å. This would indicate that only for CD=10 Å we assist to the network dismantle that completely impairs the PSI functioning. These outcomes give us information about the whole feasibility of the PSI network functioning, showing that system efficiency is negatively affected by lowering CD; even so, the possibility to perform FRET in the whole network is severely impaired only reaching the lowest value of the CD. The Forster energy transfer process (FRET) appears to have played the role of a design constraint shaping the evolution of photosynthetic organisms and have had profound influence in shaping their architecture [38,39,43] Our outcomes shed light on the structural evolution of the PSI, showing how the PSI network formed by FRET links is a resilient system with a large *‘window of feasibility”*, holding the capacity to perform the energy transfer in the whole network even when deprived of the most of FRET links.

### 4.3 Connectivity among different chromophore types

When analysing the connectivity among different types of chromophores we find that for higher value of CD there is higher LHCs-Core linkage density than intra LCHs. Only decreasing CD up to CD=50 Å, the intra-LHCs and the LHCS-Core links density become comparable (Fig. 7). In other words, lowering the CD produces a faster decrease of the LCHs-Core links with small effect in decreasing the intra-LHCs links. On one hand, this indicates that LHCs are spatially closer among them than they are with the Core Chls. On the other hand, this shows that a large amount LHCs-Core links is of lower FRET efficiency. For this reason, for CD=50 Å the network is formed by higher FRET links shared in similar number in LHCs-Core and intra-LHCs links (Fig. 7). This finding would describe a PSI system composed by a fraction of robust and highly probable energy transfer links, that we can call ‘*low regime links*”, shared roughly in the same quantity between intra-LHCs and LHCs-Core links. These links may act as baseline mechanism supporting the system functioning for below-optimum regime, such as for example when the light harvesting is lower. Differently, when the efficiency is higher (CD>50 Å), even reaching optimal values (for CD>80 Å), a large amount of LHCs-Core links becomes likely to perform the energy transfer supporting the low regime links. We call these numerous and low FRET the ‘*high regime links*’ and they may represent an internal mechanism for boosting the PSI system toward higher level of functioning.

### 4.4 The nodes-chromophores attack

The nodes removal (attack) strategies simulations have the aim to mimic the loss of functionality of the chromophores of the PSI network, giving us information on how different network nodes contribute to the functioning efficiency of the overall system [2,3]. In normal conditions when chromophores lose their functionality they get readily substituted in the turnover process [55,56]. In the case the turnover machinery is impaired the system loses chromophores making the photosystem functionality deteriorate progressively. The decrease in the number of chromophore in the photosynthesizing organisms may be the result of genetic mutations, chemical damage (such as the loss of Chls), pollution or the changing environment of the chromophores [3]. The reduction of the Chls number (or functionality) can also be associated to natural phenomena as the yellowing of senescing leaves and autumnal colour changes in in the foliage of deciduous trees, but also in seeming evergreens, such as ferns and algae [57].

Removing a node-chromophore in the PSI network would affect the system in two ways. On one hand, it removes from the system one entry gate for light, thus lowering the light harvesting capacity. On the other hand, it removes all the links relating to that node-chromophore breaking up the paths for the FRET passing into that node. The paths disruption elongates the energy transfer travel among nodes producing a decrease in the network efficiency (*Eff*) (Fig. 2).

We observed that removal of nodes corresponding to carotenoid molecules (Cars) induces lower average damage to the network functioning (*Eff* and *LCC*) than the random nodes removal (Fig. 8). In fact, Cars are nodes possessing only outgoing links and very low betwenness centrality (*BC*) not playing the role of halfway hubs for the FRET paths in the network. Thus, their removal will not induce the FRET paths disruption, resulting in a lower network efficiency decrease, outlining the small Cars importance for the structural efficiency of the energy transfer, and relegating their main role for other important functions, as broadening the spectrum of absorbable light and photoprotection [6].

Similarly, we find that random removal of Chls belonging to LHCs causes lower average damage to network functioning (*Eff* and *LCC*) (Fig. 8). As for carotenoids, LHCs nodes show low betweenness centrality (Figs 5–6), indicating their minor role for routing the shortest paths in the network and consequently the lower damage to the PSI network efficiency (*Eff*) caused by their removal. For this reason, similar to the Cars nodes the LHCs would play a major function of light entry gate as proposed by [58].

Differently, we find that the random removal of Core Chls (Rand Core) triggers the greatest decrease in the network functioning (*Eff* and *LCC*) (Fig. 8). Furthermore, the single removal of Core Chls triggers the highest decrease in the network efficiency (*Eff*) than all the other types of nodes (δ indicator in Figs 5–6). The higher damage to the system EET induced by the removal of the chlorophylls from the Core may be explained with their peculiar embedding within the structure of the PSI network. The Core Chls show the highest node betweenness centrality, unveiling them as major articulation points for the FRET shortest paths among the network nodes (Fig. 5). Their removal breaks up many higher efficiency FRET paths causing greater damage to global energy transfer efficiency, raising the Core Chls as the key network component directly involved in the increase of EET of the *P. Sativum* PSI, confirming by means of network theory previous results from experimental analysis [59,60].

## 5. Conclusion

In this paper we modelled the PSI of the *Pisum Sativum* as complex interacting network. First, we discovered that progressively removing the weaker FRET links decreases the network efficiency, while the nodes connectivity is still preserved. This finding would unveil a large window of feasibility of the PSI functioning, in which the photosynthetic process it is completely impaired only losing almost all the links in the network. In other terms, the PSI networked system would be highly resilient to the malfunctioning of FRET links. Second, we discovered that the Core Chls removal triggers the fastest functioning decrease, indicating the chromophores in the core of the PSI network as the main actor boosting the EET. Since the nodes removal simulates the chromophores malfunctioning, caused for example by the senescing leaves, biological diseases, pollution chemical damage or others [57], the PSI energy transfer efficiency would be more vulnerable to the core chlorophylls functionality loss. The outcomes presented here open new promising researches, for example building and comparing the energy transfer efficiency of the PSI network of different natural and agricultural plant species, or improving the light-harvesting mechanisms of artificial photosynthesis both in plant agriculture [59,60] and in the field of solar energy applications [61].

## Acknowledgments

MB and FS acknowledges financial support from Fondazione Cariplo, grant n° 2018-0979. This project has received funding from the European Research Council (ERC) under the European Union’s Horizon 2020 research and innovation programme (grant agreement No. [816313]). Many thanks to Prof. Guglielmo Lanzani whose comments greatly improved the first version of this paper.

