## Appendix for "Modeling the photosynthetic system I as complex interacting network"

### A.1 Nodes-chromophores indices

**Table A1:** Indices of the nodes-chromophores in the PSI network.

| | Indices |
|---|---|
| **Core** | 7, 8, 9, 10, 11, 12, 13, 14, 15, 16, 17, 18, 19, 20, 21, 22, 23, 24, 25, 26, 27, 28, 29, 30, 31, 32, 33, 34, 35, 36, 37, 38, 39, 40, 41, 42, 43, 44, 45, 46, 47, 48, 49, 51, 52, 59, 60, 61, 62, 63, 64, 65, 66, 67, 68, 69, 70, 71, 72, 73, 74, 75, 76, 77, 78, 79, 80, 81, 82, 83, 84, 85, 86, 87, 88, 89, 90, 91, 92, 93, 94, 95, 96, 98, 99, 102, 103, 104, 105, 106, 108, 113, 114, 115, 118, 119, 120 |
| **Cars** | 1, 2, 3, 4, 5, 6, 53, 54, 55, 56, 57, 58, 100, 101, 107, 109, 110, 111, 112, 116, 117 |
| **P700** | 50, 97 |
| **LHC1** | 121, 122, 123, 124, 125, 126, 127, 128, 129, 130, 131, 132, 133, 134 |
| **LHC2** | 135, 136, 137, 138, 139, 140, 141, 142, 143, 144, 145, 146, 147, 148 |
| **LHC3** | 149, 150, 151, 152, 153, 154, 155, 156, 157, 158, 159, 160, 161, 162, 163, 164 |
| **LHC4** | 165, 166, 167, 168, 169, 170, 171, 172, 173, 174, 175, 176, 177, 178, 179 |

### A.2 PSI network properties

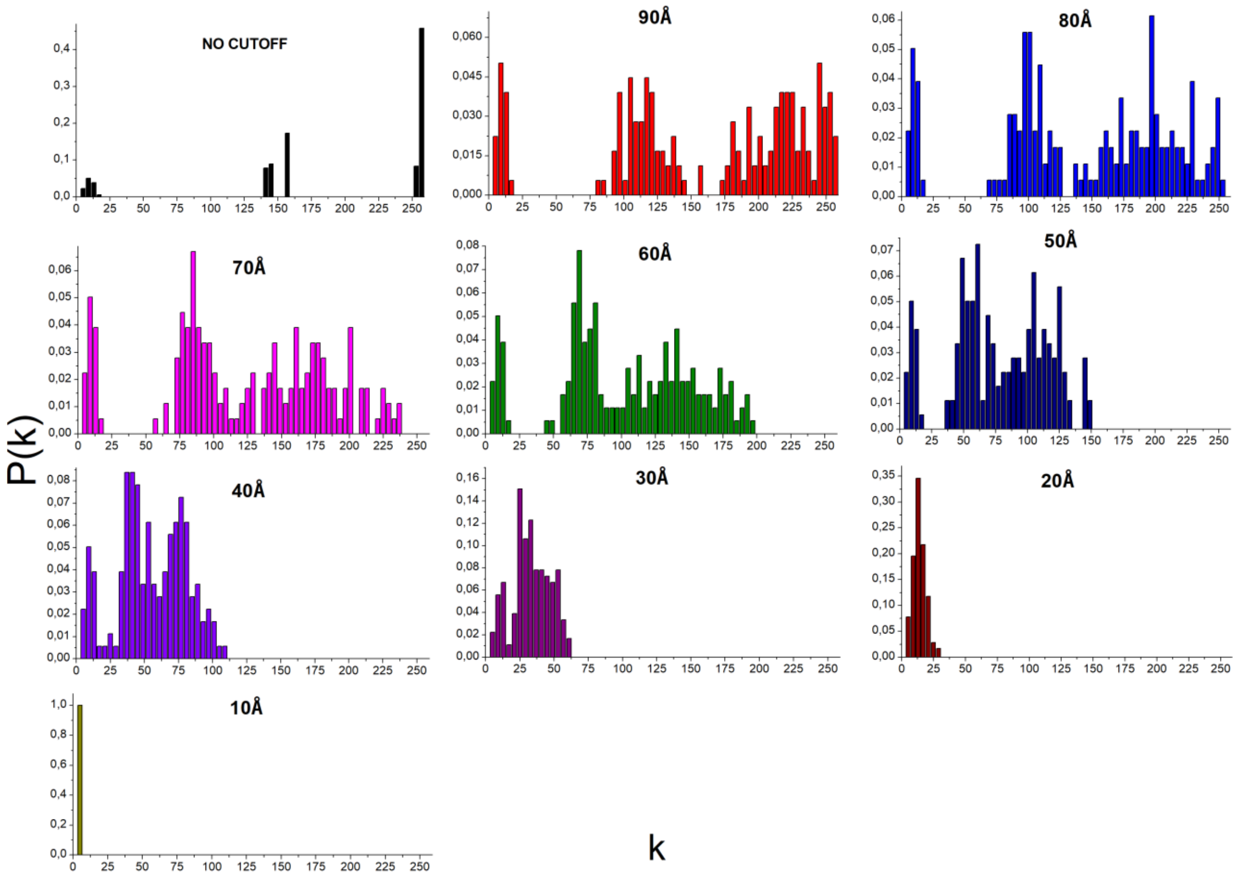


**Figure A1:** Degree distribution *P(k)*, depending on the cut-off distance (CD).

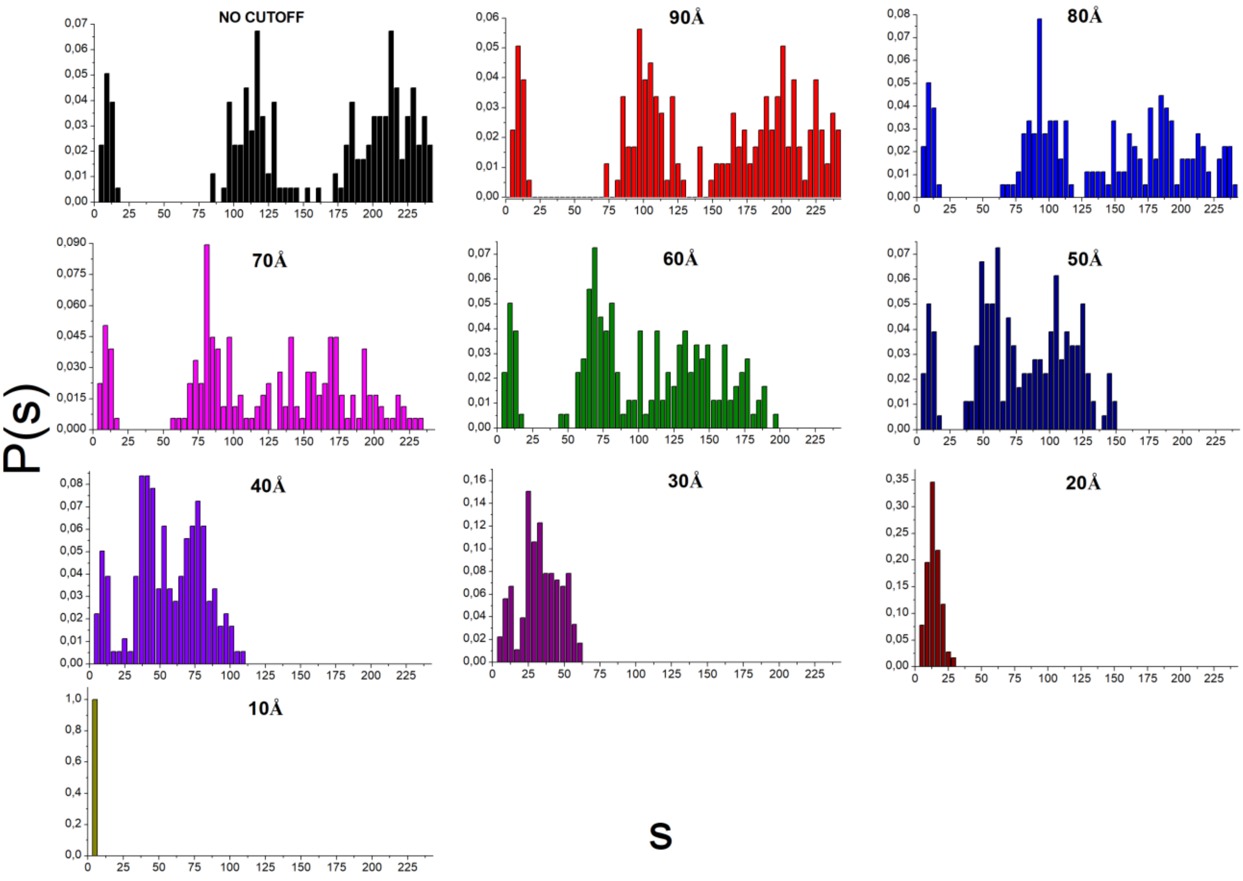


**Figure A2**: Strength distribution *P(s)* depending on the cut-off distance (CD).

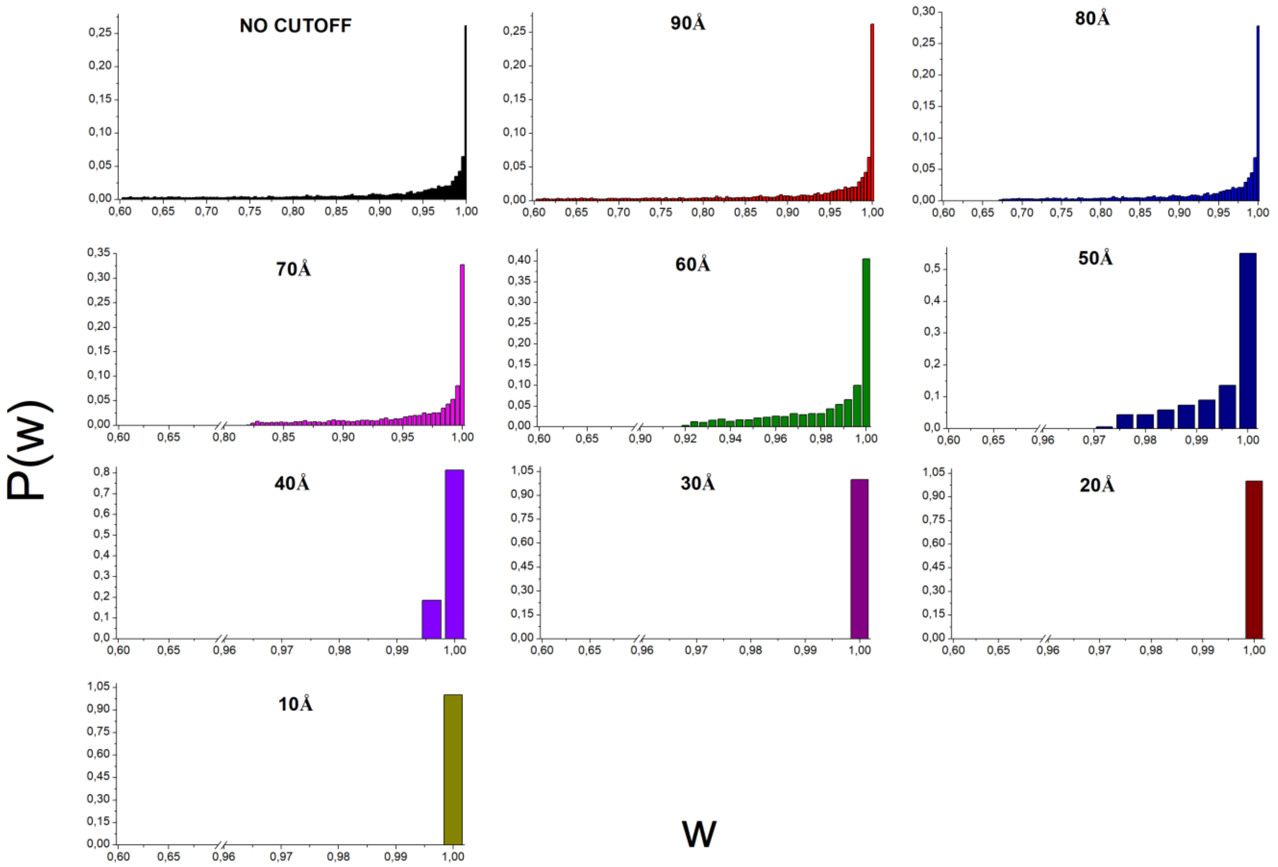


**Figure A3**: Weights distributions *P(w)* of the edges, depending on the cut-off distance (CD).

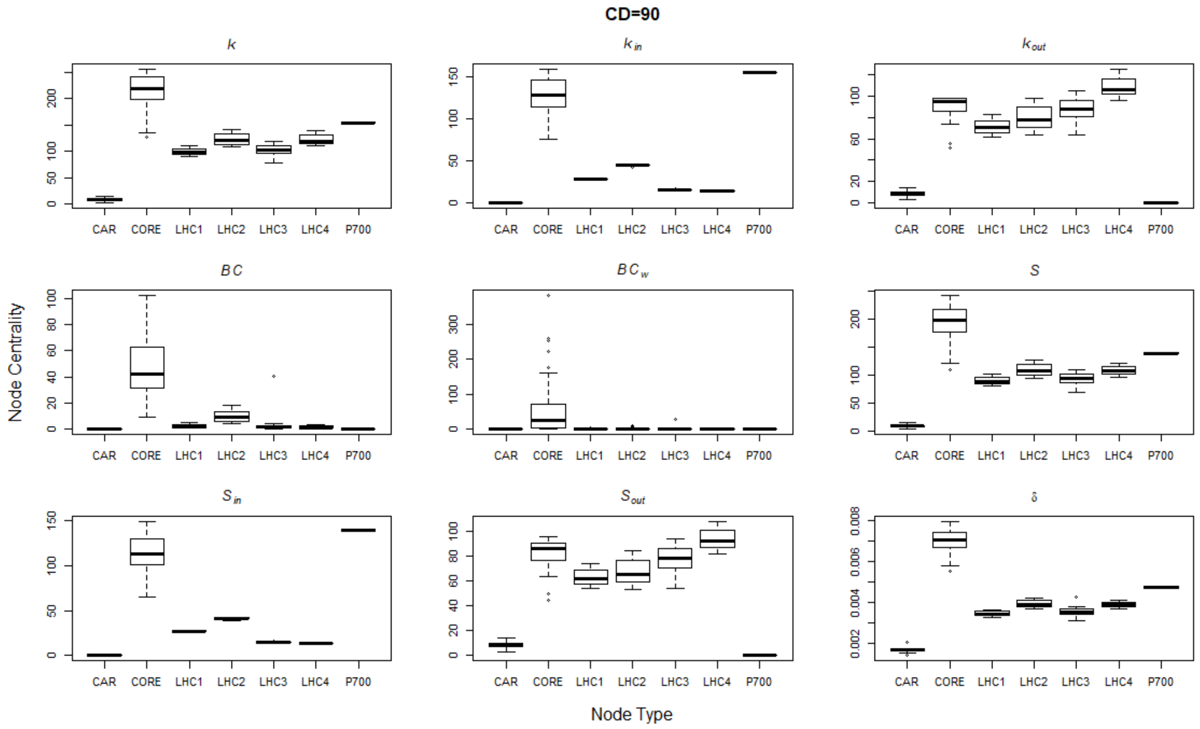


**Figure A4**: Node types *vs* nodes centrality feature for the PSI network features CD=90 Å. Keys are: $k$ degree of the nodes, $k_{out}$ out degree, $k_{in}$ in-degree; $BC$ binary directed betwennness centrality; $BCw$ binary directed betwennness centrality; $S$ node strength, $S_{in}$ ingoing node strength, $S_{out}$ outgoing node strength; δ decrease in the directed network efficiency after single node removal.

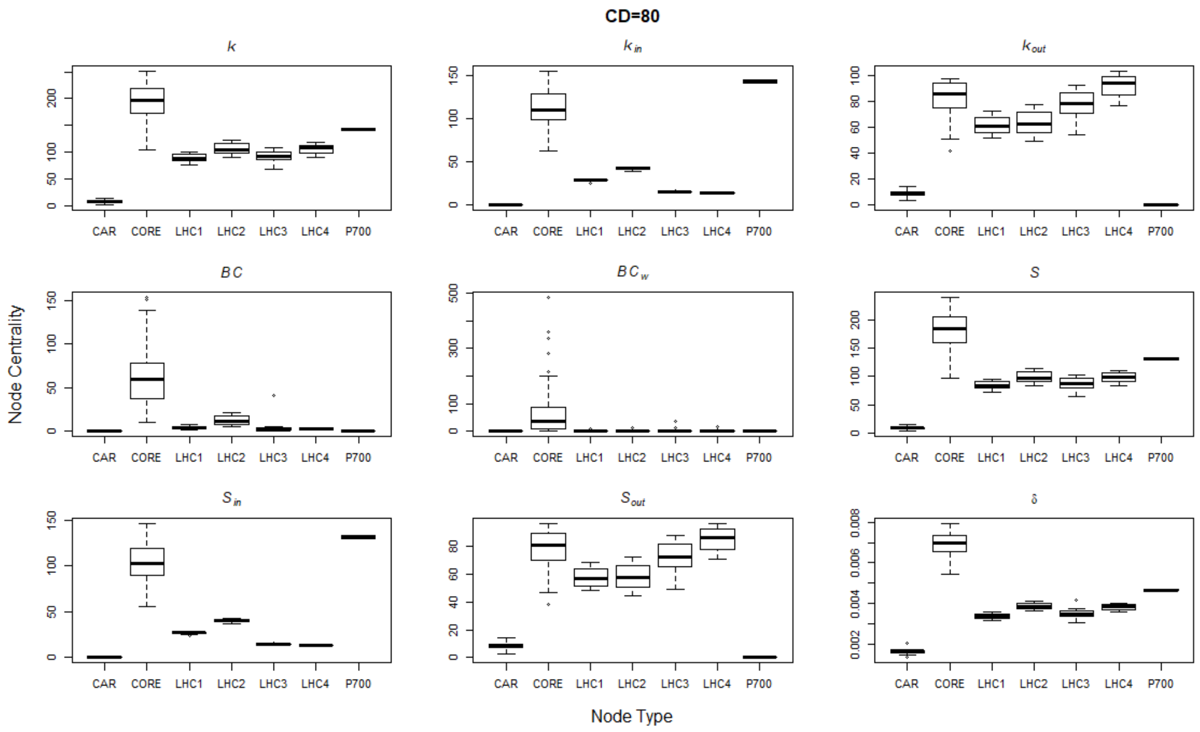


**Figure A5**: Node types *vs* nodes centrality feature for the PSI network features for CD=80 Å. Keys are: $k$ degree of the nodes, $k_{out}$ out degree, $k_{in}$ in-degree; $BC$ binary directed betwennness centrality; $BCw$ binary directed betwennness centrality; $S$ node strength, $S_{in}$ ingoing node strength, $S_{out}$ outgoing node strength; δ decrease in the directed network efficiency after single node removal.

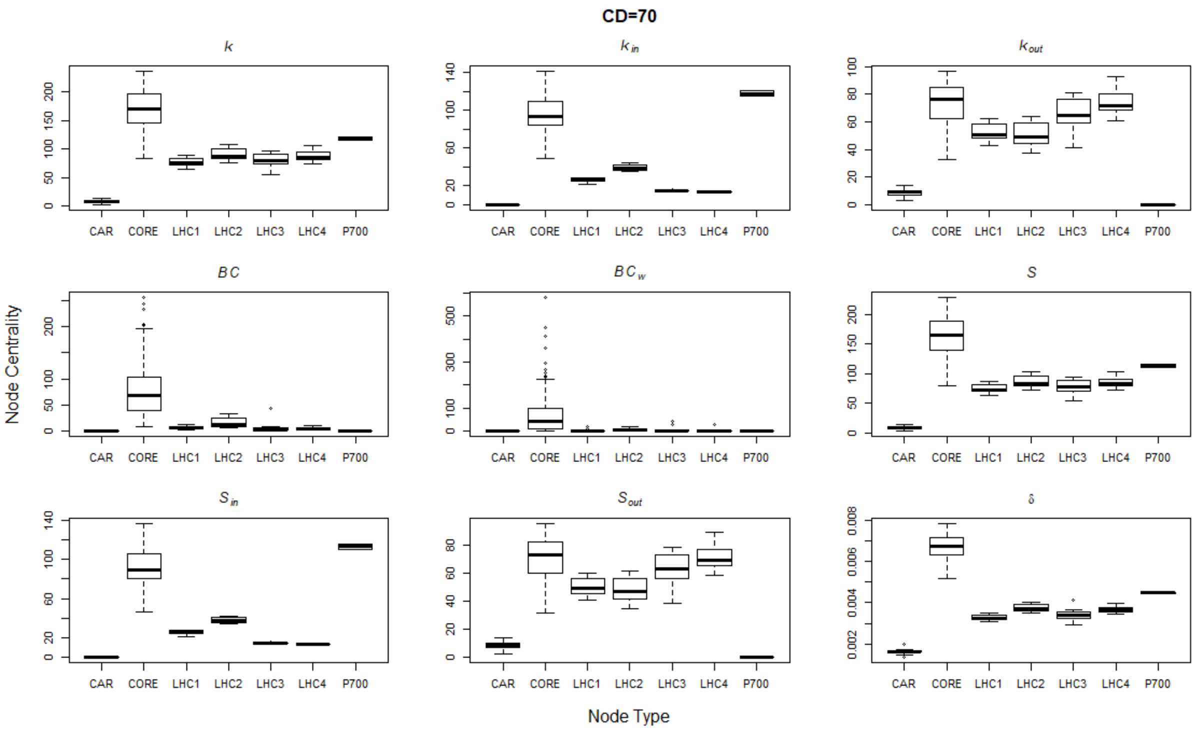


**Figure A6**: Node types *vs* nodes centrality feature for the PSI network features for CD=70 Å. Keys are: $k$ degree of the nodes, $k_{out}$ out degree, $k_{in}$ in-degree; *BC* binary directed betwennness centrality; *BCw* binary directed betwennness centrality; $S$ node strength, $S_{in}$ ingoing node strength, $S_{out}$ outgoing node strength; δ decrease in the directed network efficiency after single node removal.

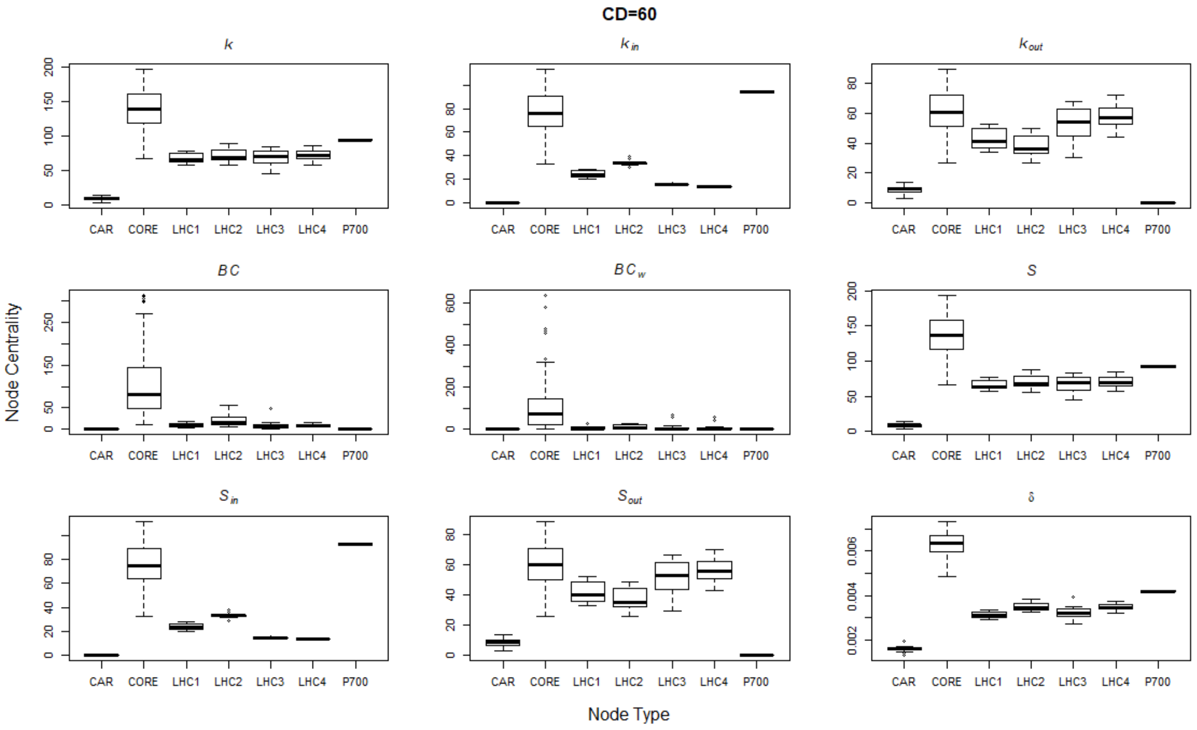


**Figure A7**: Node types *vs* nodes centrality feature for the PSI network features for CD=60 Å. Keys are: *k* degree of the nodes, $k_{out}$ out degree, $k_{in}$ in-degree; *BC* binary directed betwennness centrality; *BCw* binary directed betwennness centrality; *S* node strength, $S_{in}$ ingoing node strength, $S_{out}$ outgoing node strength; δ decrease in the directed network efficiency after single node removal.

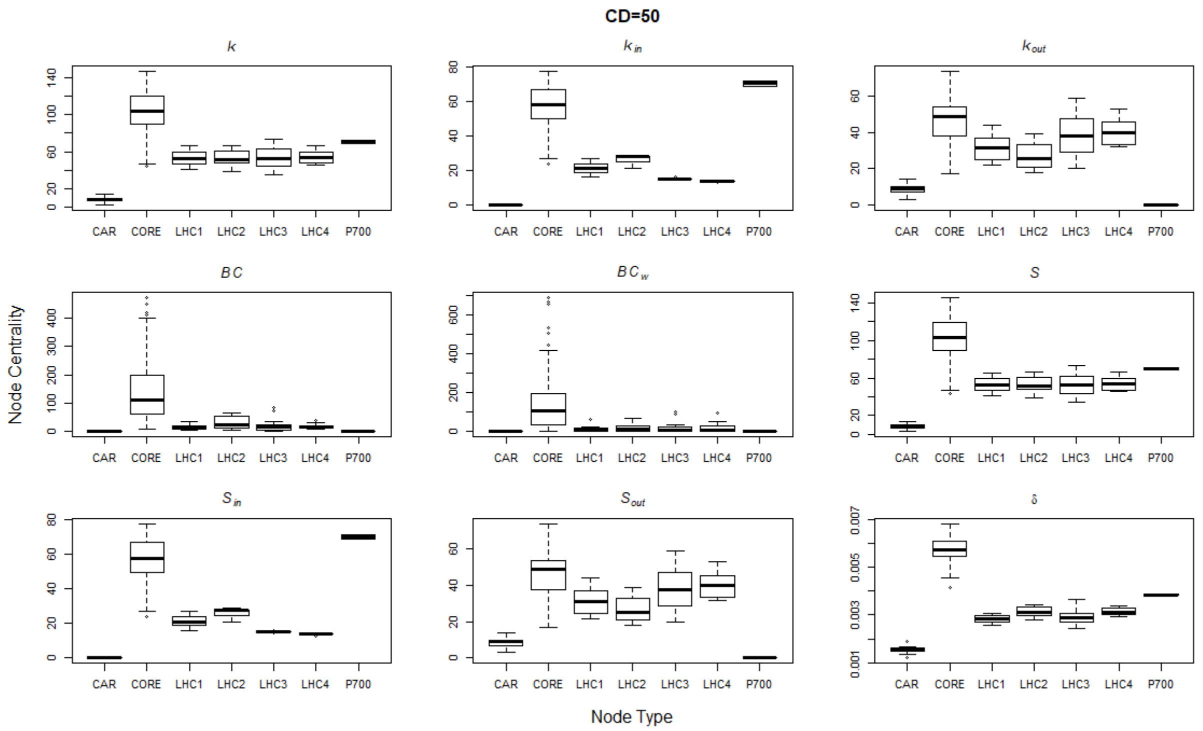


**Figure A8**: Node types *vs* nodes centrality feature for the PSI network features for CD=50 Å. Keys are: $k$ degree of the nodes, $k_{out}$ out degree, $k_{in}$ in-degree; *BC* binary directed betwennness centrality; *BCw* binary directed betwennness centrality; $S$ node strength, $S_{in}$ ingoing node strength, $S_{out}$ outgoing node strength; δ decrease in the directed network efficiency after single node removal.

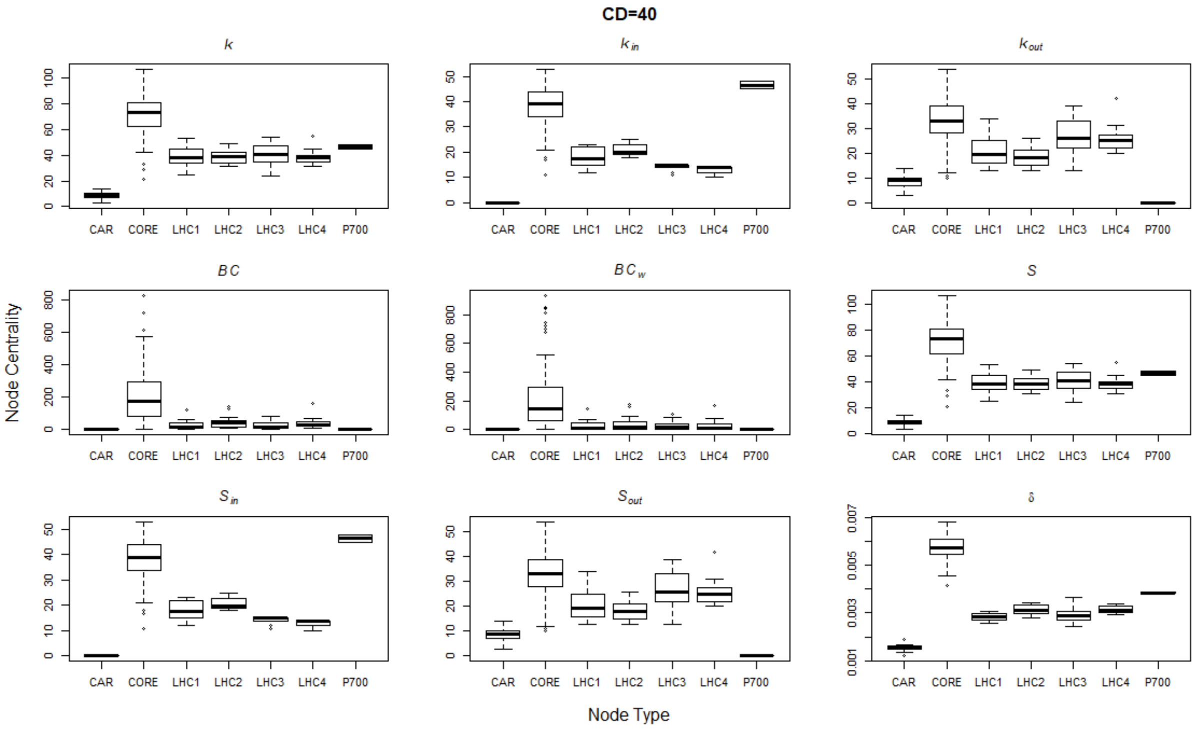


**Figure A9**: Node types vs nodes centrality feature for the PSI network features for CD= 40 Å. Keys are: *k* degree of the nodes, $k_{out}$ out degree, $k_{in}$ in-degree; *BC* binary directed betwennness centrality; *BCw* binary directed betwennness centrality; *S* node strength, $S_{in}$ ingoing node strength, $S_{out}$ outgoing node strength; δ decrease in the directed network efficiency after single node removal.

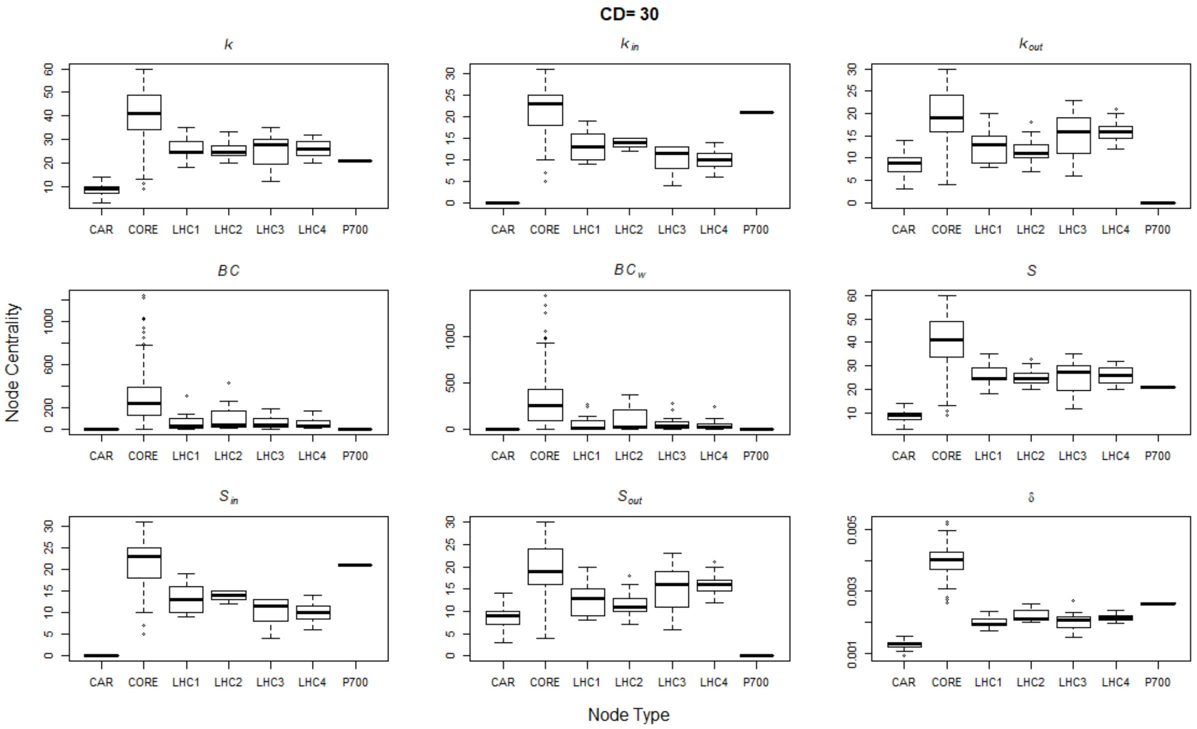


**Figure A10**: Node types *vs* nodes centrality feature for the PSI network features for CD= 30 Å. Keys are: $k$ degree of the nodes, $k_{out}$ out degree, $k_{in}$ in-degree; *BC* binary directed betwennness centrality; *BCw* binary directed betwennness centrality; $S$ node strength, $S_{in}$ ingoing node strength, $S_{out}$ outgoing node strength; δ decrease in the directed network efficiency after single node removal.

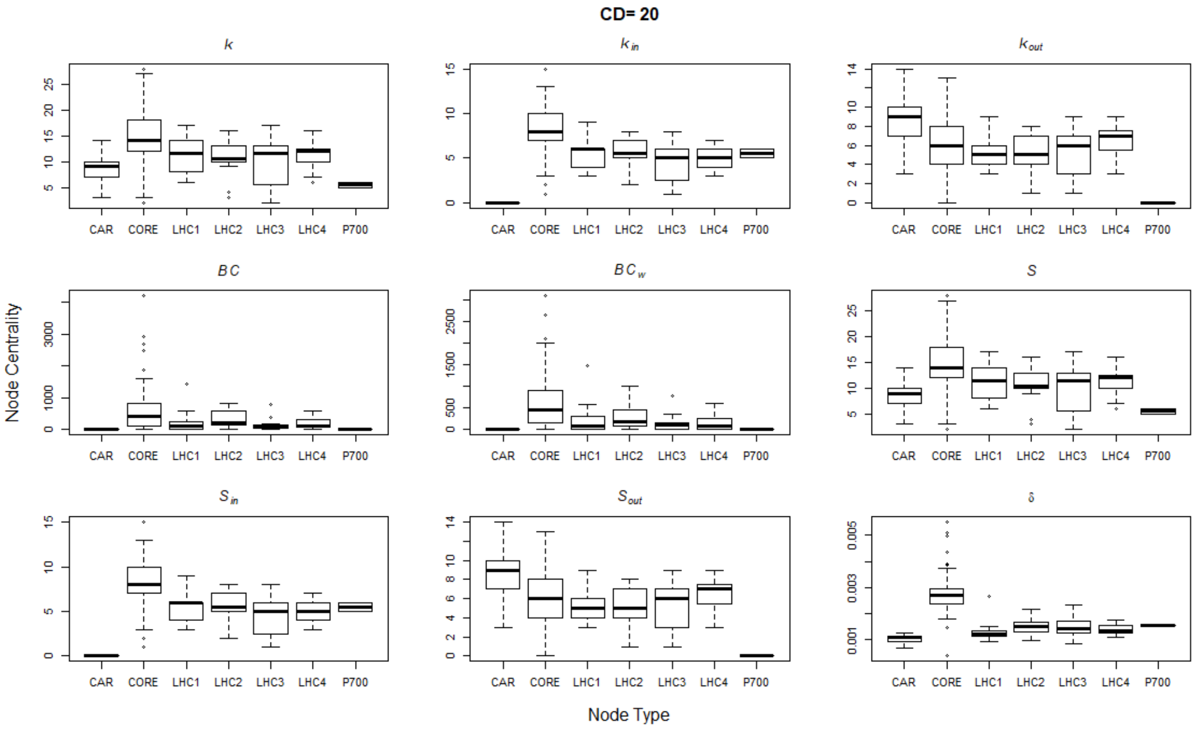


**Figure A11**: Node types *vs* nodes centrality feature for the PSI network features for CD= 20 Å. Keys are: *k* degree of the nodes, $k_{out}$ out degree, $k_{in}$ in-degree; *BC* binary directed betwennness centrality; *BCw* binary directed betwennness centrality; *S* node strength, $S_{in}$ ingoing node strength, $S_{out}$ outgoing node strength; δ decrease in the directed network efficiency after single node removal.

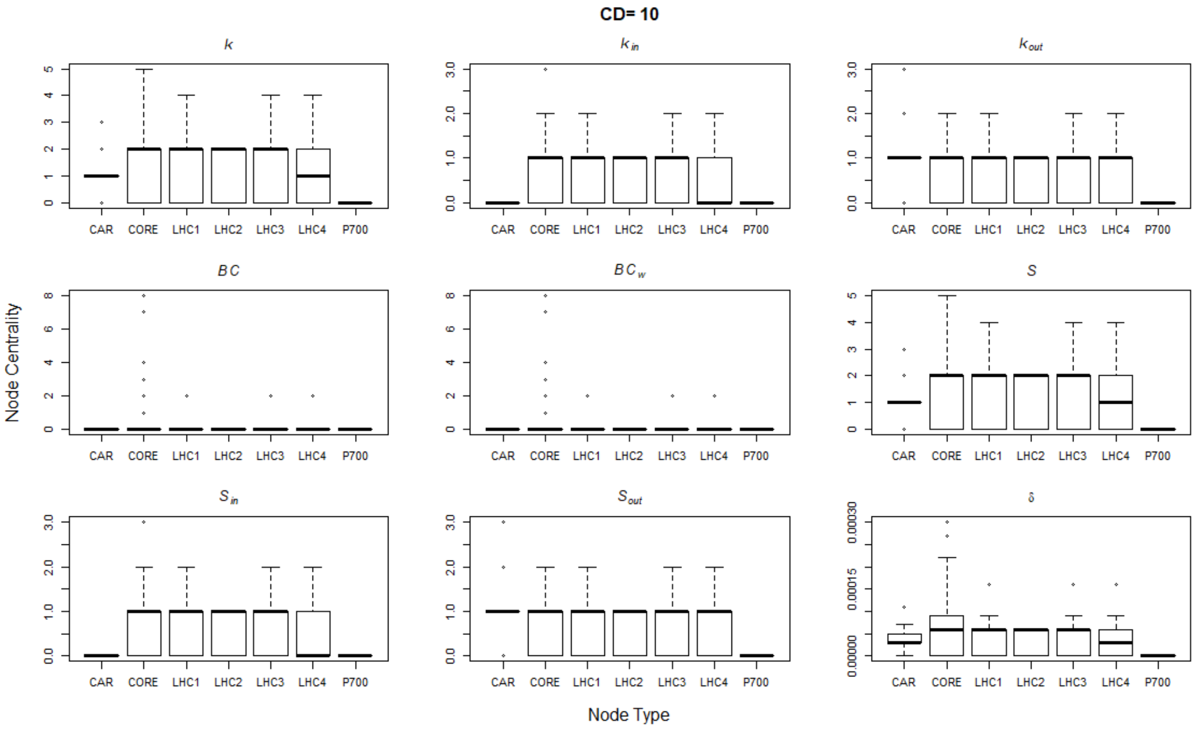


**Figure A12**: Node types *vs* nodes centrality feature for the PSI network features for CD= 10 Å. Keys are: $k$ degree of the nodes, $k_{out}$ out degree, $k_{in}$ in-degree; *BC* binary directed betwennness centrality; *BCw* binary directed betwennness centrality; $S$ node strength, $S_{in}$ ingoing node strength, $S_{out}$ outgoing node strength; δ decrease in the directed network efficiency after single node removal.

### A.3 Nodes Removal simulations outcomes

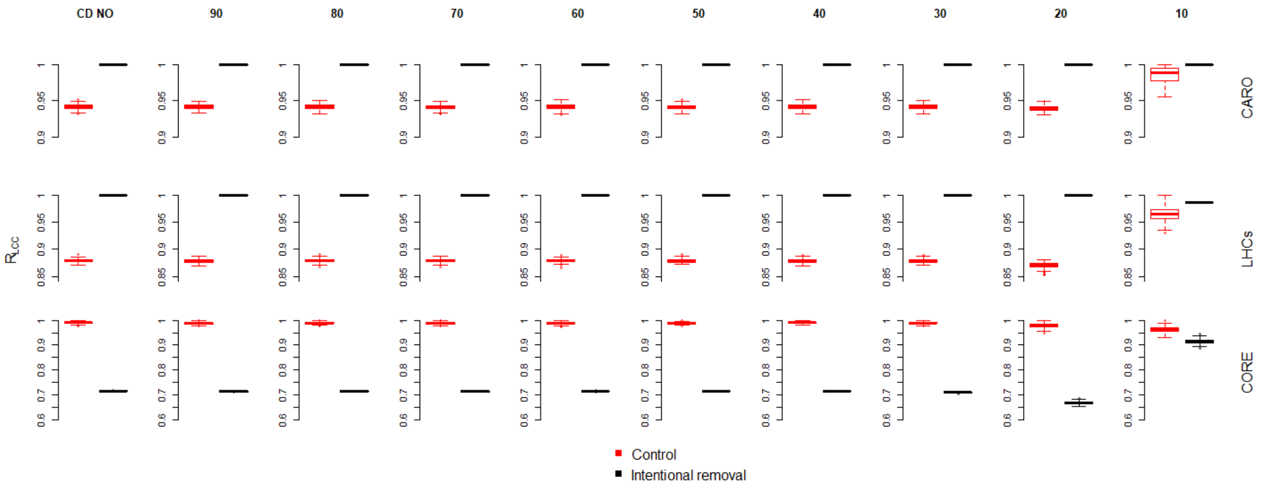


**Figure A13**: PSI network robustness ($R_{LCC}$) under the random removal (Rand) and intentional random nodes removal strategies for the different CD. The damage in the network during the nodes removal is evaluated with the weakly largest connected component $LCC_{weak}$. The intentional random removals: top row carotenoids, half row LHCs and bottom row Core Chls.
